# Conserved Lysine in transmembrane helix 5 is key for the inner gating of the LAT transporter BasC

**DOI:** 10.1101/2024.03.26.586791

**Authors:** Joana Fort, Adrià Nicolàs-Aragó, Luca Maggi, Maria Martinez Molledo, Despoina Kapiki, Niels Zijlstra, Susanna Bodoy, Els Pardon, Jan Steyaert, Oscar Llorca, Modesto Orozco, Thorben Cordes, Manuel Palacín

**Author notes:** contributed equally (Joana Fort, Adrià Nicolàs-Aragó).

## Abstract

L-amino acid transporters (LATs) play a key role in a wide range of physiological processes. Defects in LATs can lead to neurological disorders and aminoacidurias, while the overexpression of these transporters is related to cancer. BasC is a bacterial LAT transporter with an APC fold. In this study, to monitor the cytoplasmic motion of BasC, we developed a smFRET assay that can characterize the conformational states of the intracellular gate in solution at room temperature. Based on combined biochemical and biophysical data and molecular dynamics simulations, we propose a model in which the conserved lysine residue in TM5 supports TM1a to explore both open and closed states within the cytoplasmic gate under apo conditions. This equilibrium can be altered by substrates, mutation of conserved lysine 154 in TM5, or transport-blocking nanobodies. Overall, these findings provide insights into the transport mechanism of BasC and highlight the significance of the lysine residue in TM5 in the cytoplasmic gating of LATs.

## Introduction

L-amino acid transporters (LATs) (SLC7A5-11 and SLC7A13 in humans) play a key role in the cellular uptake and distribution of amino acids. LATs are thus relevant for a wide range of physiological processes, including protein synthesis, signal transduction, and the maintenance of cellular homeostasis. Defects in LATs lead to a range of disorders, including aminoacidurias and neurological disorders (Fotiadis *et al*, 2013). In this regard, mutations in SLC7A7 and SLC7A9 cause the rare aminoacidurias Lysinuric protein intolerance (LPI) (Torrents *et al*, 1999) and cystinuria (Feliubadaló *et al*, 1999), respectively. Similarly, mutations in SLC7A8 are associated with age-related hearing loss (Espino Guarch *et al*, 2018) and cataracts (Knöpfel *et al*, 2019), and the first mutation in SLC7A10 has recently been described to cause the neurological disorder hyperekplexia (Drehmann *et al*, 2023). Moreover SLC7A5, SLC7A8, and SLC7A11 overexpression has been observed in many types of cancer (Kanai, 2022; Xiong *et al*, 2019; Koppula *et al*, 2021), where these transporters confer a metabolic advantage for tumour growth and progression. These examples underscore that a fundamental understanding of LAT transport mechanisms is highly relevant not only for understanding physiology but also for clinical applications.

LATs feature the so-called APC fold, which is characterized by a 5+5 inverted repeat of transmembrane helices (TM1 to TM5 and TM6 to TM10 in LATs, Fig 1A). At the heart of the inverted repeat lie TM1 and TM6, each with an unwound segment, housing the main binding site of the substrates (del Alamo *et al*, 2022), dividing their transmembrane helices into two parts (TM1a-TM1b and TM6a-TM6b from the N- to C-terminus). These half-TMs are part of cytoplasmic and extracellular gates, especially TM1a on the cytoplasmic side and TM6a on the extracellular side.

**Figure 1.**
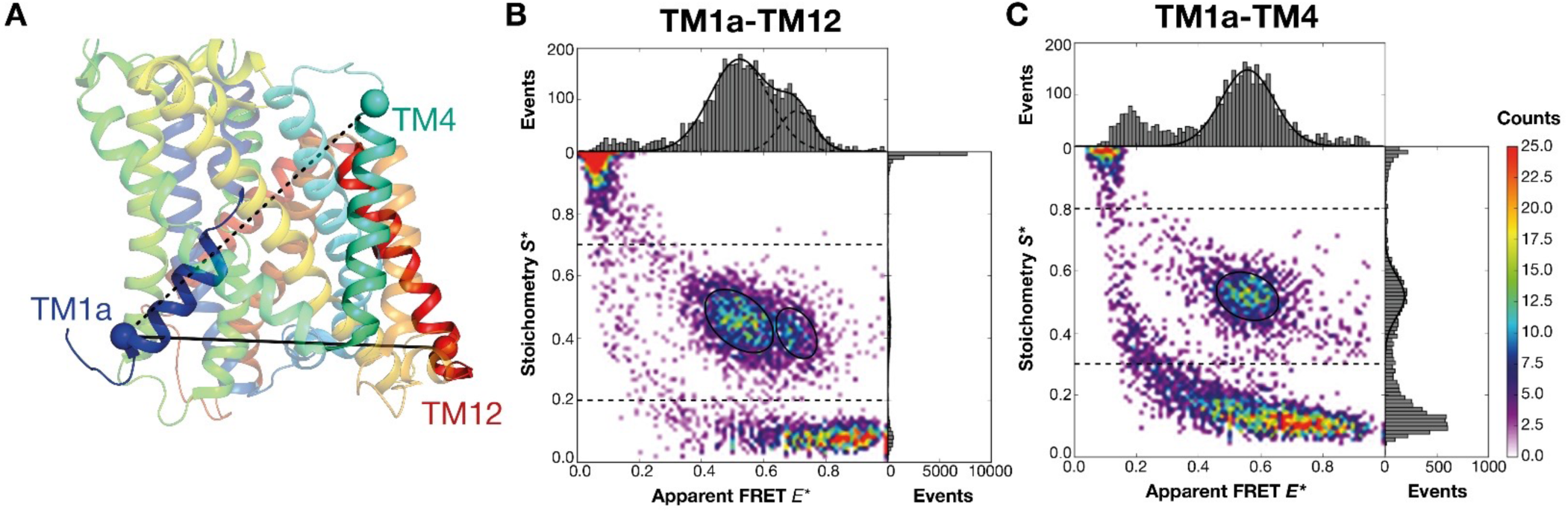
Schematic of smFRET assays and representative data for BasC. **A.** Residue positions used for dye labelling in smFRET experiments using the inward-open conformation of BasC (PDB ID 6F2W). Ball symbols represent the C-α atoms of Ile7 in TM1a, Thr120 in TM4, and Cys427 in TM12. Dashed and solid grey lines represent the double-cysteine variants studied, TM1a-TM4 and TM1a-TM12, respectively. **B, C.** Two-dimensional *E\** vs *S\** histograms from ALEX-FRET of BasC double-cysteine variants TM1a-12 (B) and TM1a-4 (C) under apo conditions with 0.06 % DDM. One-dimensional histograms show projections of the data within limited S-range of 0.3-0.8 for 1D-E* and the full projection for S*. The distributions were fitted with a model using one or two gaussian functions showing mean values of 0.52 (σ = 0.09) and 0.71 (σ = 0.06) for TM1a-TM12 (B) and of 0.56 (σ = 0.09) for TM1a-TM4 (C). Thresholding and data analysis procedures are described in detail in the Material and Methods section.

In recent years, structural studies have been complemented by biophysical techniques that can monitor changes in conformational dynamics even at room temperature in solution. Single-molecule Förster Resonance Energy Transfer (smFRET) (Ha *et al*, 1996; Lerner *et al*, 2018; Hellenkamp *et al*, 2018; Lerner *et al*, 2021; Agam *et al*, 2023) for example, has been successfully used to study the dynamics of membrane transporters at the molecular level (Bartels *et al*, 2021; Gouridis *et al*, 2015, 2015; Dyla *et al*, 2017; Husada *et al*, 2018; Yang *et al*, 2018; de Boer *et al*, 2019; Lasitza-Male *et al*, 2020; Debruycker *et al*, 2020). Relevant to this work, smFRET could help to show that the LeuT transporter, a bacterial homolog of the neurotransmitter sodium symporter (NSS) family with APC fold(Yamashita *et al*, 2005), undergoes a rocking motion and switch from an outward-facing to an inward-facing conformation (Zhao *et al*, 2010, 2011; Tavoulari *et al*, 2016; Malinauskaite *et al*, 2014; Terry *et al*, 2018; Khan *et al*, 2020). The proposed rocking motion of the bundle domain (TM1, TM2, TM6 and TM7) versus the scaffold domain (TM3, TM4, TM8 and TM9) between outward-facing to inward-facing conformations is consistent with the general mechanism observed in other members of the APC transporter superfamily, including LAT structures (Errasti-Murugarren *et al*, 2019; Yan *et al*, 2019; Lee *et al*, 2019; Rodriguez *et al*, 2021; Parker *et al*, 2021; Yan *et al*, 2020a; Wu *et al*, 2020; Lee *et al*, 2022; Jeckelmann *et al*, 2022; Yan *et al*, 2020b, 2021; Lee *et al*, 2023).

Despite the same fold and some mechanistic aspects, the NSS and LAT families of transporters do not share significant sequence identity; moreover, they have distinct transport mechanisms. NSS transporters are sodium symporters (Rudnick *et al*, 2014), whereas LATs are amino acid exchangers (Fotiadis *et al*, 2013) that transit between inward and outward-facing conformations with substrate bound in the binding site. Furthermore, human LATs and the bacterial LAT BasC show lower Km for substrates in the outward (μM range) than in the inward (mM range) conformations (Meier *et al*, 2002; Bartoccioni *et al*, 2019). This property allows for the harmonization of substrate concentrations on both sites of the cell membrane (Gauthier-Coles *et al*, 2021, 2023). Interestingly, in BasC and human LATs, the side chains of two conserved residues, a tyrosine in TM7 (Tyr236) and a lysine in TM5 (Lys154 in BasC), occupy similar sites as the two sodium atoms in NSS transporters (Na1 and Na2 sites, respectively) and both residues are important for the asymmetric affinity for the substrate (Errasti-Murugarren *et al*, 2019). Moreover, Na2 in LeuT is known to be involved in the opening and closing of the inner gate (Krishnamurthy *et al*, 2009; Stolzenberg *et al*, 2017). Interestingly, the conserved lysine residue in TM5 is key for transport activity in LATs. Interestingly, the homologous lysine in SLC7A7 is mutated to glutamate (Lys194Glu) in a patient with LPI (Sperandeo *et al*, 2008). Moreover, the mutation of the homologous lysine to glutamate or to alanine in BasC and SLC7A10 results in a dramatic loss of function (Errasti-Murugarren *et al*, 2019). Whether this conserved lysine residue plays a role in the inner gating of LATs is unknown.

In the present work, to monitor cytoplasmic movement of BasC, we developed a smFRET assay that can characterize the conformational states of the ligand-free (apo) and ligand-bound (holo) form. The assay uses fluorophore labelling of TM1a-TM12 and TM1a-TM4 (Fig 1A) to track the motion of TM1a. Using specific nanobodies and point mutation of the conserved lysine in TM5 reveal a key role of this residue for gating at the cytoplasmic side of BasC. Based on the combined data and molecular dynamics simulations, we propose a model in which the conserved lysine residue in TM5 supports TM1a to explore both open and closed states within the cytoplasmic gate under apo conditions. This equilibrium can be altered by the substrate addition, mutation of the TM5 lysine, or incubation with nanobodies (Nbs). Overall, these findings provide insights into the transport mechanism of BasC and highlight the significance of the lysine residue in TM5 in the cytoplasmic gating of LATs.

## Results

### BasC samples different conformations under apo conditions

To monitor the movement of the inner gate of BasC in solution at room temperature, we developed a smFRET assay that tracks its conformational changes. To this end, two distinct double-cysteine variants of BasC were designed. Residue Ile7 in TM1a was present in both (Fig 1A), because TM1a is predicted to move strongly during the closing of the inner gate in LATs. The equivalent position to Ile7 in BasC is Leu53 in human LAT SLC7A5, which differs by 11.5 Å between the inward-open (Protein Data Bank, PDB ID 6IRS) (Yan *et al*, 2019) and outward occluded structures (PDB ID 7DSQ) (Yan *et al*, 2021). Moreover, smFRET studies of LeuT seeking to track the inner gate movement used a similar position in TM1a (Zhao *et al*, 2010; Tavoulari *et al*, 2016; Terry *et al*, 2018; Khan *et al*, 2020). In BasC, Ile7 in TM1a was combined with two other residues in the protein: Thr120 in TM4, forming the TM1a-TM4 pair, and Cys427, the only natural cysteine in BasC, in TM12, forming the TM1a-TM12 pair (Fig 1A). Both BasC variants with cysteine-residues in TM1a-TM4 (mutations Ile7Cys-Thr120Cys-Cys427Ala) and TM1a-TM12 (mutation Ile7Cys) retained approximately 60% and 80% activity, respectively, in proteoliposomes (PLs) (Fig EV1).

We used maleimide-conjugated FRET fluorophores to stochastically label the cysteine pairs in the two protein variants. Fluorophore-labelled proteins solubilized in detergent DDM were then studied in free diffusion using an in house-built confocal microscope (Gebhardt *et al*, 2021). Fig 1B and C show representative data of BasC for TM1a-TM12 (Fig 1B, dye pair Alexa 546 / Alexa 647) and TM1a–TM4 (Fig 1C, dye pair SCy3 / SCy5). We used alternating laser excitation, µsALEX (Kapanidis *et al*, 2004; Hohlbein *et al*, 2014), to derive information on both the apparent FRET efficiency E* and brightness ratio S* (Fig 1B/C). The two-dimensional histogram allowed us to focus our analysis on donor-acceptor-labelled BasC by selecting an S* region between 0.3 and 0.8, indicated by a dashed rectangle.

In the TM1a-TM12 pair, we observed a bimodal distribution with two distinct E*-populations (Fig 1B). Fitting of the distribution with two gaussian functions gave mean E* values of 0.52 (σ=0.09) and 0.71 (σ=0.06) for both states (Fig 1B). The TM1a-TM4 pair showed only a single peak with a mean E* value of 0.56 (σ=0.09) (Fig 1C). These results under ligand-free apo conditions suggest that the protein can adopt at least two distinct conformations (Fig 1B). The observation of the single peak of the TM1a-TM4 pair (Fig 1C) is attributed to a reduced sensitivity of the sCy3-sCy5 pair, since the connection axis of the residues in TM1a-TM4a lies not directly within the direction of movement of TM1a. Additional evidence for this interpretation comes from the Cy3-sCy5 labelling of the TM1a-TM12 pair, where also a broader distribution is observed, yet no two discrete populations (Fig EV3), showing the higher sensitivity of the Alexa dyes in the assay.

### Substrate addition induces a closing movement of TM1a

In the next step, we used the smFRET assay to study the effect of the substrates L-alanine and L-serine and the competitive inhibitor L-glutamine (Fig EV2) on BasC. Upon addition of L-alanine, we observed an increase in the high-FRET population and a slight shift in the low-FRET population for TM1a-TM12 labelling (Fig 2A). Quantitatively, the substrate increased the high-FRET fraction, where TM1a and TM12 were closer, from 25% to 45% approximately (Fig 2B). Under similar conditions, the peak observed for TM1a-TM4 shifted towards higher E* values upon addition of both L-alanine and L-serine (Fig 2C). In contrast, the addition of L-glutamine did not cause a shift (Fig 2C). To obtain statistics from independent experiments, we normalized the ligand-induced E* shifts and calculated ΔE* as the difference between E* in the presence or absence of ligand (Appendix Fig S1). Fig 2D shows the observed peak shifts (ι1E*) of TM1a-TM4, confirming a statistical increase ι1E*>0 for substrates but no change in ι1E*≈0 for the inhibitor. The increase observed in E* indicates a decrease in the interprobe distances between the labelled positions in TM1a and TM4, consistent with a closing movement of the inner gate. Indeed, positions TM1a-TM12 labelled with SCy3 and SCy5 showed a similar tendency when substrate was added (Fig EV3A).

**Figure 2.**
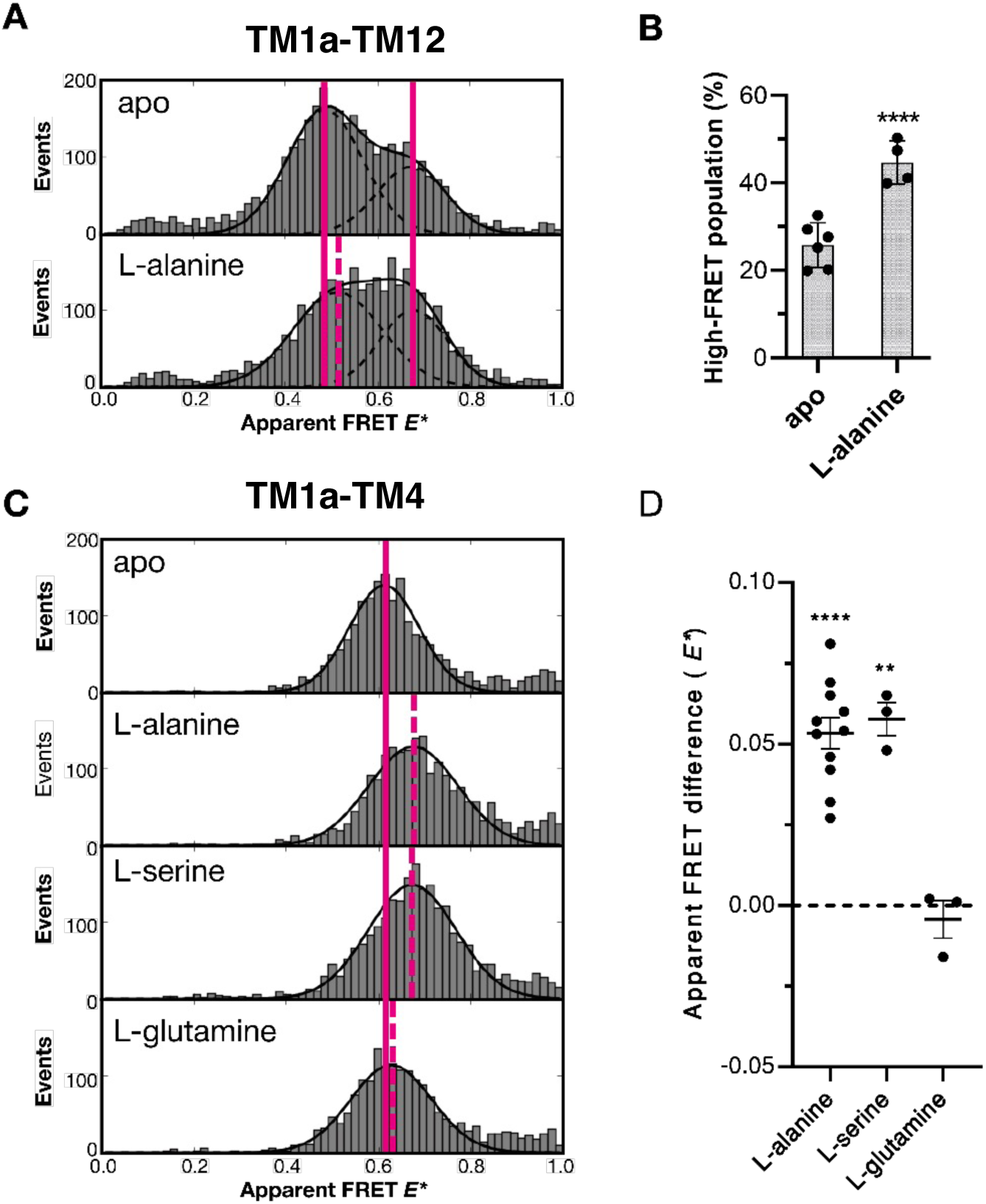
The impact of substrates on the conformational state of TM1a. smFRET data of BasC variant TM1a-12 labelled with Alexa 546 and Alexa 647 **(A and B)** and TM1a-TM4 variant labelled with sCy3-sCy5 **(C and D)** in the presence and absence of different amino acids. **A, C.** Representative histograms of apparent FRET *E\** distributions for TM1a-TM12 and TM1a-TM4 variants under different buffer conditions. Vertical solid pink lines represent the mean E* value of the apo state and dashed lines that of the respective amino acid. **B.** Quantitative analysis of the population distribution between low and high FRET for apo and holo conditions in TM1a-TM12. We statistically compared the relative abundance of both states in independent repeats (black dots) by plotting mean ± standard error of the mean (SEM) of the percentage of high-FRET state. Paired T-tests were conducted against apo measurements from the same day. P-values are indicated by asterisks (**** < 0.001). **D.** Quantitative analysis of apparent FRET *E\** differences between holo and apo state of BasC TM1a-TM4. For this, mean values of the holo distribution were statistically compared with an apo measurement from the same day (ΔE*, black dots). Mean ΔE* ± SEM are displayed and paired T-tests were performed using the absolute *E\** values comparing apo and holo measurements of the same day. P-values are denoted by asterisks (**** < 0.001 and ** < 0.01).

In addition, we measured the substrate-induced movement of the TM1a-TM4 pair under increasing concentrations of L-alanine. The response showed saturation >100mM with an estimated EC50 around 40 mM (Fig EV3B). This value is very similar to the reported dissociation constant Kd of BasC (∼35 mM), determined by thermostability-based substrate binding assays (Errasti-Murugarren *et al*, 2019). The smFRET results support the notion that substrate binding to BasC favours a closer conformation TM1a.

### Substrate-induced movement is prevented by specifically blocking TM1a

To confirm the involvement of TM1a in the substrate-induced movement detected by smFRET, we used three specific nanobodies (Nbs) targeting the cytoplasmic side of BasC: Nb53, Nb71, and Nb74. These Nbs were selected on the basis of their capacity to inhibit the efflux of 10 µM L-alanine, as shown in Fig EV4A, following the protocol previously used for Nb74 (Errasti-Murugarren *et al*, 2019). Briefly, proteoliposomes (PLs) with randomly inserted BasC are externally treated with a saturating concentration of blocking Nbs. Nbs targeting the cytoplasmic side can interact with the inside-out oriented BasC molecules, thereby inhibiting the efflux of L-alanine at 10 µM, as demonstrated for Nb53, Nb71, or Nb74 (Fig EV4). In contrast, Nbs targeting the extracellular side of BasC interacts with right-side up BasC molecules and, consequently, inhibit the influx of L-alanine at 10 µM, as shown for Nb58 (Fig EV4).

The incubation of BasC with Nb71 resulted in the disappearance of the high FRET population for the TM1a-TM12 pair (Fig 3A) and diminished ι1E*-effects of L-alanine in the TM1a-TM4 pair (Fig 3B). Interestingly, the incubation of BasC with Nb53 did not affect the ratio of the two peaks in the BasC TM1a-TM12 pair (Fig 3A) or alter the effects of L-alanine in the TM1a-TM4 pair (Fig 3B). Also, the incubation of BasC with Nb74 did not diminish the L-alanine-induced movement of the intracellular gate (Fig EV4B).

**Figure 3.**
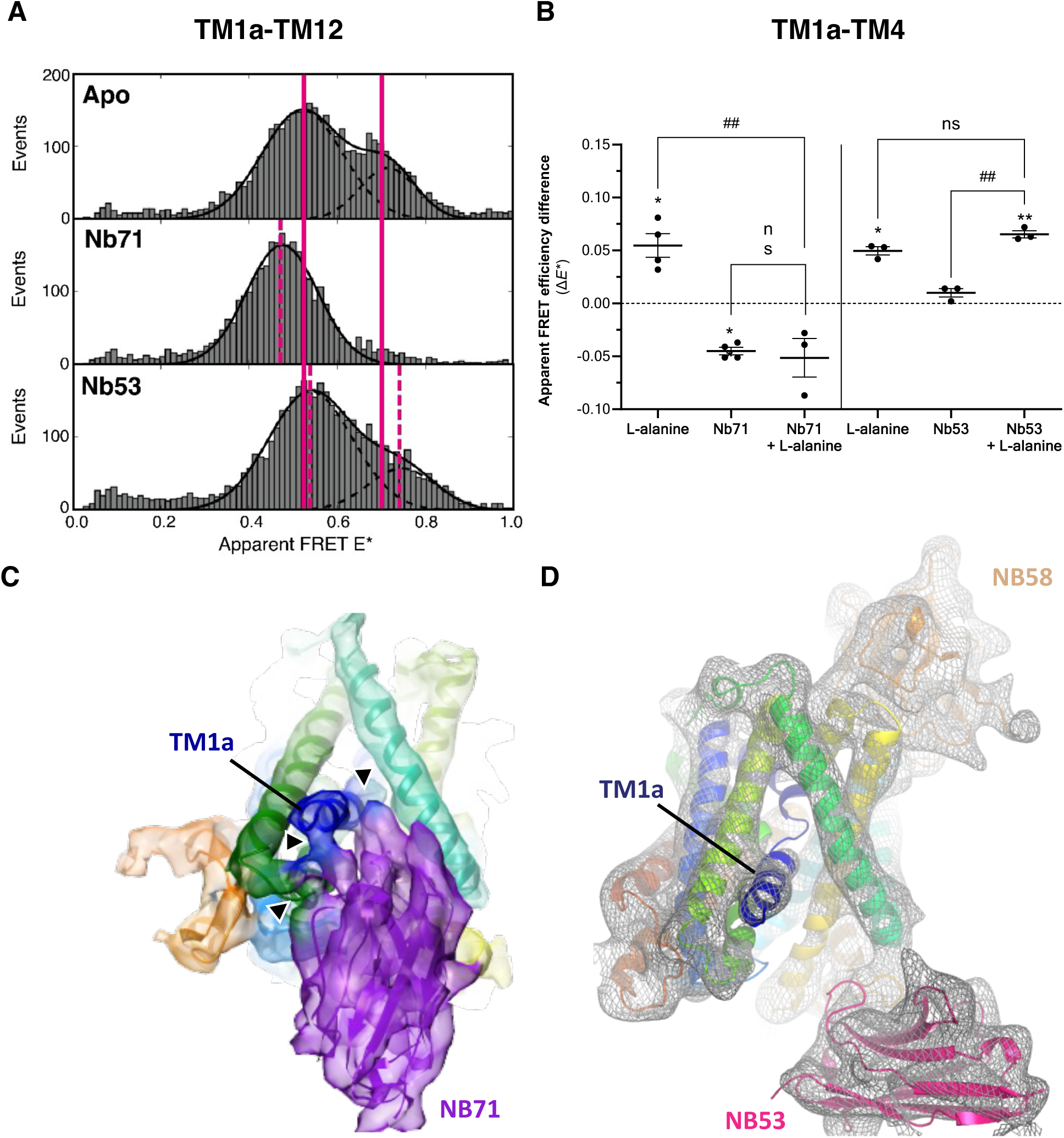
The effect of nanobodies on conformational states BasC. **A.** Comparison of E* distribution for the TM1a-12 variant, in the absence and presence of 1 µM of the indicated Nanobodies. Vertical pink lines represent mean E* values for the apo untreated populations, while dashed red lines indicate mean E* values for populations incubated with each respective Nb indicating slight deviations. **B.** Differences in ΔE* are observed in the TM1a-4 variant (labelled with Alexa 546 and Alexa 647) when incubated with substrate (300 mM) in the presence or absence of the indicated Nbs (1 µM). Mean ± SEM values are represented. Statistical significance was determined using paired T-tests comparing each condition against the apo (denoted by asterisks) or L-alanine (L-Ala) (denoted by hashes) conditions of the same day, with p-values * < 0.05 and **, ## < 0.01 or ns (no statistical significance). **C.** Low-Resolution Cryo-EM volume (∼6.0 Å) of Nb71-BasC with fitted BasC model from PDBID 6F2G (in rainbow) and a Nb71 homology model (in purple). The depicted volume shows direct interactions between Nb71 and TM1a and TM6 of BasC which is shown by arrows. **D.** Low-Resolution X-ray map (∼6.0 Å, grey mesh) of the Nb53-BasC-Nb58 complex showing no direct interaction of Nb53 with TM1a of BasC. Models shown in rainbow cartoon of BasC from PDBID 6F2G, Nb53 model (pink) and Nb58 model (orange) have been slightly refined inside the X-ray map to show interacting domains. The blocking effect of Nb58 in the influx via BasC in PLs (Fig. EV4A) supports the extracellular interaction of this nanobody with BasC.

To explain these smFRET results on a molecular level, we conducted low-resolution structural studies of BasC. Cryo-EM was used for a characterization of BasC complexed with Nb71 (Appendix Fig S2, Table S2) and crystallography for BasC complexed with Nb53, alongside with Nb58 to increase crystal contacts. The obtained low-resolution structures show that Nb71 was able to establish direct interactions with both TM1a and TM6b (Fig 3C), providing an explanation for the complete reduction of movement of the inner gate. Conversely, Nb53 showed no interactions with TM1a or the rest of the bundle domain (Fig 3D) —observations consistent with its lack of impact on the L-alanine-induced movement of TM1a. As previously reported, Nb74 did not interact with TM1a but did with TM6b in the bundle domain (PDB ID 6F2G, Errasti-Murugarren *et al*, 2019), thereby pointing to the closing of TM1a independently of TM6b.

The structural information obtained was thus crucial for understanding the observed behaviour of the three Nbs in smFRET experiments and confirm that the L-alanine-induced shift in smFRET experiments corresponds to the closing of TM1a, as part of the cytoplasmic gate. These results strongly support the notion that the substrate-induced changes in FRET efficiency E* and the distributions reflect the occlusion of TM1a when the substrate interacts to the binding site. Indeed, specific blocking of TM1a with Nb71 prevented this movement.

### The fully conserved lysine residue in TM5 (Lys154) is needed for the closing of TM1a

The lysine residue in TM5, i.e., Lys154 in BasC, is a conserved residue among LATs that is described as critical in the transport mechanism (Errasti-Murugarren *et al*, 2019). To elucidate the role of Lys154 in the substrate-induced movement of TM1a in BasC, we conducted additional experiments using a Lys154Ala variant (Fig 4). smFRET experiments using the TM1a-TM12 and TM1a-TM4 pairs were performed within the Lys154Ala BasC variant. Interestingly, the two distinct populations observed in the TM1a-TM12 pair in wild-type (WT) BasC were not detected in the presence of Lys154Ala (K154A) under apo conditions or after substrate addition (Fig 4A and B). Moreover, the addition of L-alanine failed to induce any shift in E* for the TM1a-TM4 pair when the Lys154Ala mutation was present (Fig 4C and D). These results reveal that Lys154 plays a crucial role in TM1a: (i) allowing it to sample different conformational states of the inner gate even in the absence of substrate and (ii) mediating the closing of the inner gate in BasC upon substrate binding.

**Figure 4.**
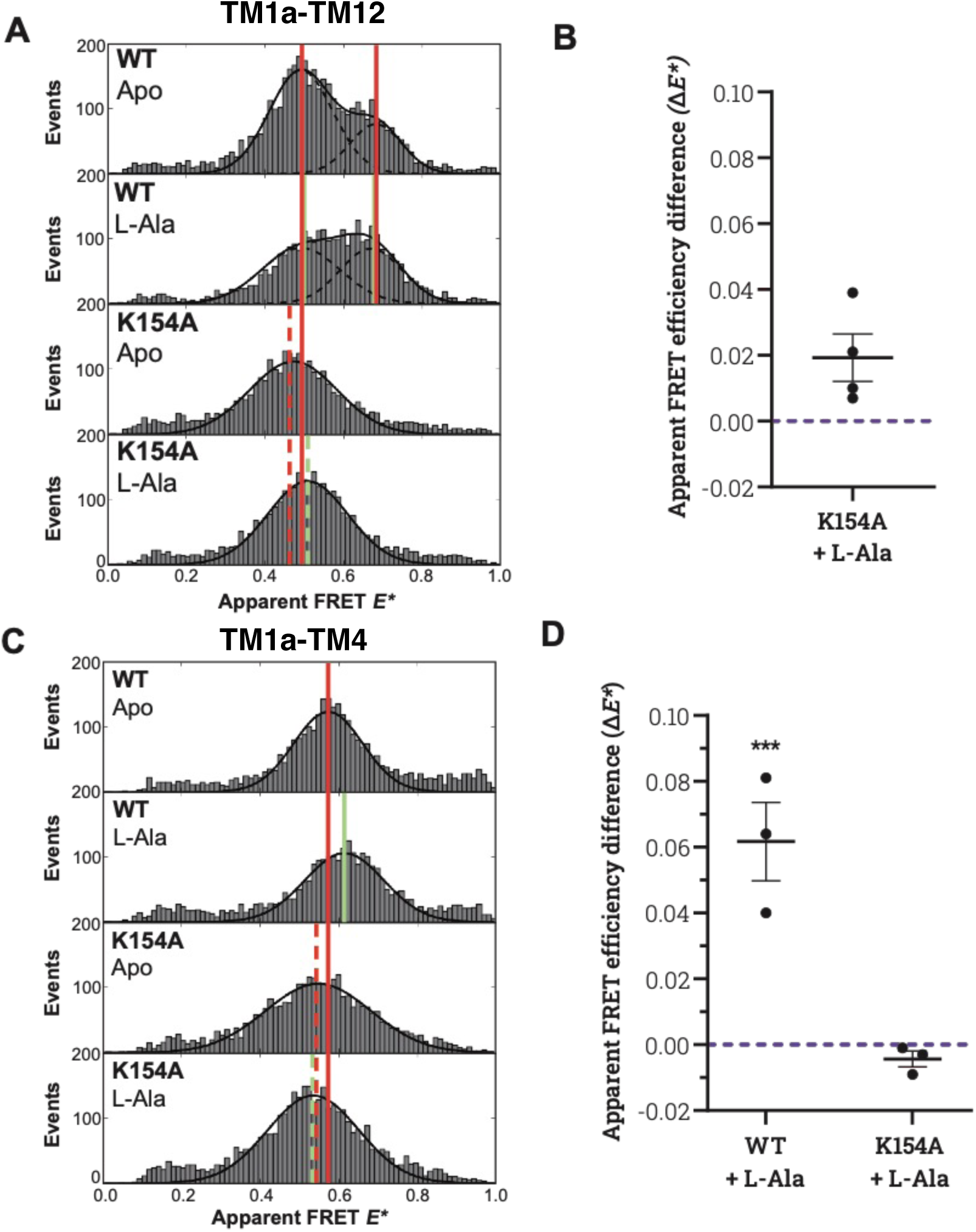
Effect of the Lys154Ala mutation on the conformational states of TM1a. **A, C.** Representative smFRET histograms illustrating the influence of the Lys154Ala (K154A) mutation on conformational states and changes for BasC double-cysteine variants TM1a-TM12 (A) and TM1a-TM4 (C). **B, D.** Apparent FRET differences Δ*E\** derived from n = 4 and n = 3 experiments on TM1a-12 (B) and TM1a-4 (D), respectively. These experiments quantify substrate-induced smFRET changes on Lys154Ala (K154A) mutant variants *versus* wild-type BasC.

### Lys154 stabilizes the unwound segment of TM1a

To allow further interpretation of our findings, we performed atomistic molecular dynamics (MD) simulations (Fig 5) that revealed how the Lys154Ala mutation destabilizes the substrate-binding site by increasing the distance between the unwound segments of TM1 and TM6 in apo conditions (Fig 5A and B). This effect was attenuated in the holo structure (Fig 5C), but remained insufficient to allow binding of the substrate in BasC. Indeed, we observed a reduction in the strength of hydrogen bonds between the substrate 2-amino isobutyrate (2AIB) and the residues belonging to the binding site (Fig 5D) located on TM1 and TM6 as described in the solved inward-facing conformation bound to 2AIB (Errasti-Murugarren *et al*, 2019). In particular, TM1 residues were more affected by this reduction, as shown by the decrease in the percentage of hydrogen bond occupancies shown in Fig 5D. As a result, the substrate was released from the binding site in two of the three replicas of the Lys154Ala mutant, whereas it remained within the binding site in the three trajectories for WT BasC (Fig EV5B). These findings support the notion that Lys154 stabilizes the unwound segment of TM1, allowing proper binding of the substrate and, probably, permitting the next step of the transport mechanism: the closing of the inner gate.

**Figure 5.**
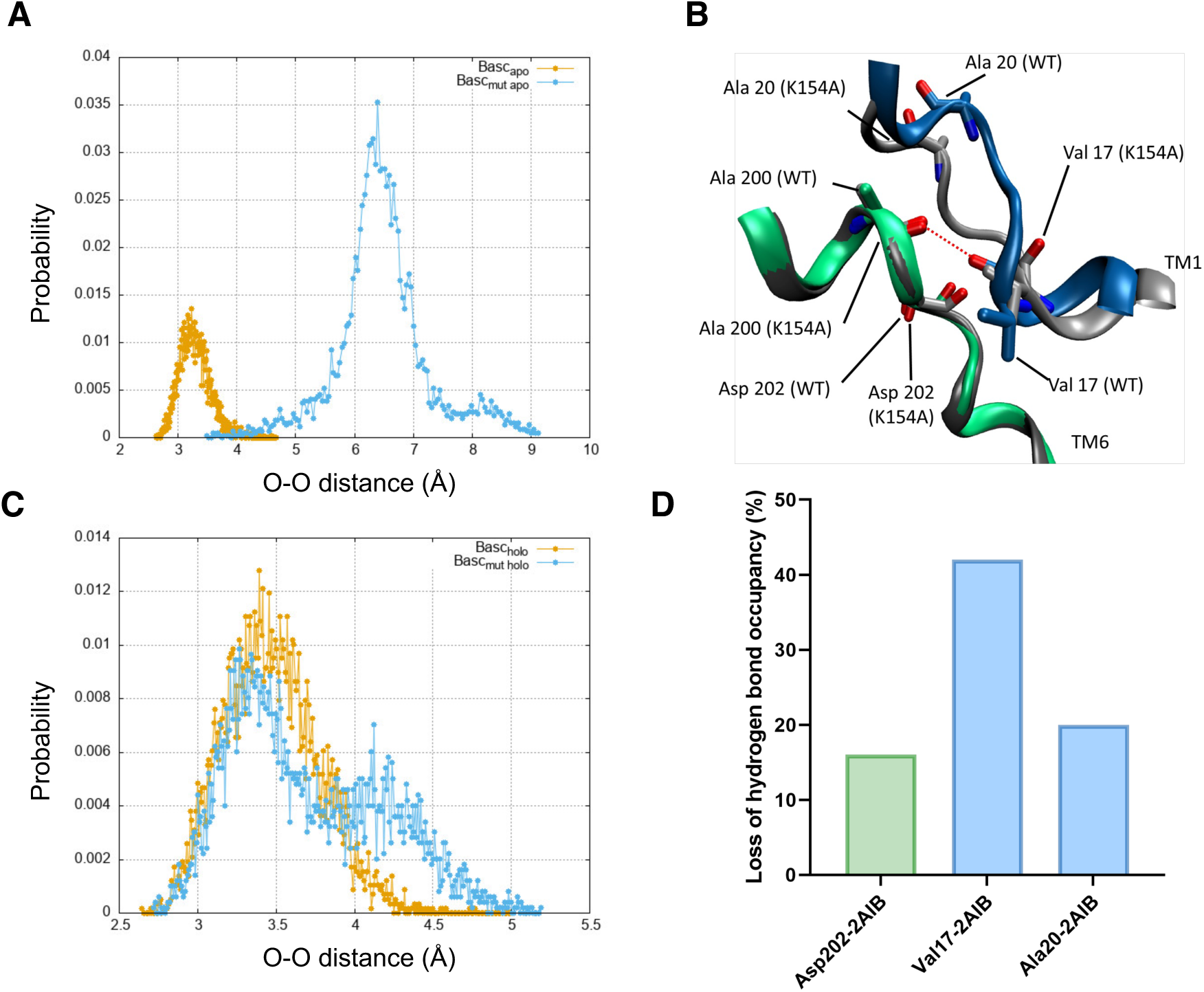
Molecular dynamics simulations of BasC. **A.** Distance distribution calculated from three replicas between the two carbonyl oxygens of Val 17 and Ala 200 in the wild type (WT) (BasC) and Lys154Ala mutant (Bascmut) of BasC in apo conditions. The graph provides insights into the structural dynamics of the binding site, emphasizing the impact of the Lys154Ala mutation on the distances between specific residues. **B.** Representative snapshot of the apo structures (BasC WT and Bascmut Lys154Ala) of the binding site extracted from molecular dynamics simulations. In the WT, TM1 and TM6 are highlighted in blue and green, respectively, while in the Lys154Ala mutation, they are both shown in grey. This visualization offers a comparative view of the binding site conformations in the presence and absence of the K154 residue. **C.** Distance distribution (in Å) calculated from three different replicas between the two carbonyl oxygens of Val 17 and Ala 200 in the WT and Lys154Ala mutant in the presence of substrate within the binding site. These data illustrate the impact of substrate binding on the structural dynamics of the binding site and how the Lys154Ala mutation influences these interactions. **D.** Percentage of loss of relative hydrogen bond occupancies for three residues displaying the strongest interaction with the substrate (2-AIB) between the WT and Lys154Ala mutation. The two blue columns represent residues from TM1, Val 17 (23.8% hydrogen bond occupancy in the WT) and Ala 20 (61% hydrogen bond occupancy in the WT). The green column corresponds to a TM6 residue, Asp 202, which shows a 33.7% hydrogen bond occupancy in the WT. These values provide insights into the disruption of hydrogen bonding patterns in the presence of the Lys154Ala mutation, highlighting its impact on substrate interactions and the stability of the binding site.

## Discussion

Based on our combined data set and MD simulations, we propose that the conserved lysine residue in TM5 helps TM1a to explore both open and closed states within the cytoplasmic gate under apo conditions, leading to an equilibrium with an open:closed ratio of approximately 3:1, which can be altered by substrates, mutation of the conserved lysine in TM5, or transport-blocking Nbs. The increased population of the closed state upon substrate addition is consistent with smFRET studies of the gating of the immobilized symporter LeuT, where transitions between the two states occur dynamically (Zhao *et al*, 2011). L-alanine substrate shifted the TM1a gating equilibrium towards the closed state in BasC (approximately 1:1 ratio of open to closed), whereas the LeuT substrate sodium pushes the equilibrium even more towards the closed state (ratio of 1:2 of open to closed) and dramatically reduces state transitions (Zhao *et al*, 2011). The gating of TM1a reflects similarities and differences between LeuT and BasC. Both transporters share the APC protein fold (Yamashita *et al*, 2005; Errasti-Murugarren *et al*, 2019), but they use distinct transport mechanisms. LeuT, as a bacterial model of Neurotransmitter:Na^+^ symporters (NSS) is an amino acid:Na^+^ symporter, whereas BasC, as a bacterial model of LATs, is an amino acid exchanger.

In LeuT, the binding of sodium ions closes the cytoplasmic gate, stabilizes the outward-facing conformation (Krishnamurthy *et al*, 2009) and facilitates the binding of the transported substrate (e.g., L-leucine) to trigger conformational transitions towards the occluded conformation. Then, partial unwinding of TM5 allows the dissociation of Na2, thereby promoting the opening of the cytoplasmic gate for the release of L-leucine into the cytoplasm (Shi *et al*, 2008). In BasC, the lateral chain of Lys154, with ε-amino group occupying a similar position to Na2 in LeuT, is involved in establishing the dynamic equilibrium of open and closed states of TM1a. This equilibrium allows sampling of the closed state by the substrate (e.g., L-alanine), which is necessary to trigger amino acid efflux from the cell, and here we establish that the highly conserved lysine residue in TM5 is central to this mechanism in LATs. Indeed, Lys154Ala mutation in BasC significantly reduces V_max_ to <10% and increases the external K_m_ value for L-alanine 10-fold, without altering the cytoplasmic K_m_ value (Errasti-Murugarren *et al*, 2019). These findings suggest that Lys154 plays a crucial role in a rate-limiting step in the BasC transport cycle, sustaining transport V_max_.

Consistent with this notion, Lys154Ala mutation freeze TM1a and the substrate is not able to stabilize the higher smFRET E* population, emphasizing the indispensability of Lys154 in mediating TM1a gating. The structural significance of Lys154 in BasC is mirrored in analogous roles within related transporters with an APC fold. The side chain of the LAT-conserved Lys residue in TM5 forms hydrogen bonds with TM1a residues, i.e., the carbonyl group of Gly15 in BasC, in all solved LAT structures (Errasti-Murugarren *et al*, 2019; Yan *et al*, 2019; Lee *et al*, 2019; Rodriguez *et al*, 2021; Parker *et al*, 2021; Yan *et al*, 2020a; Wu *et al*, 2020; Lee *et al*, 2022; Jeckelmann *et al*, 2022; Yan *et al*, 2020b, 2021). Moreover, in transporter structures with closed cytoplasmic gates, *e. g.*, human LAT1 and the related bacterial Cationic amino acid Transporter GkApcT, the conserved TM5 Lys residue bridges TM1a and TM8 (Jungnickel *et al*, 2018; Yan *et al*, 2021), thereby pointing to a pivotal TM1a-TM5-TM8 interaction for cytoplasmic gate occlusion. Similarly, in NSS transporters, coordination bonds of Na2 link residues in TM1a, TM5 and TM8, stabilizing the outward-facing conformation (Malinauskaite *et al*, 2014; Stolzenberg *et al*, 2017). This parallelism underscores the significance of Lys154 in orchestrating the inner gating of BasC. However, while this might explain the decreased L-alanine transport V_max_, it is counterintuitive that defective closing of the inner gate would increase the extracellular K_m_ of the Lys154Ala mutant.

Interestingly, our MD simulations of the substrate-bound inward-facing BasC conformation, although limited in capturing the tilting of TM1a (Appendix Fig S3), provided valuable insights. During these simulations, removal of the hydrogen-bonding capacity of the Lys154 side chain (Lys154Ala mutant) led to changes in the conformation of the unwound segment of TM1, which is a key component of the substrate-binding site in BasC and mammalian LATs in inward- and outward-facing conformations (Errasti-Murugarren *et al*, 2019; Yan *et al*, 2019, 2020a; Rodriguez *et al*, 2021; Yan *et al*, 2021; Lee *et al*, 2023). Consequently, the Lys154Ala mutation weakens interaction with the substrate and leads to the release of the substrate to the cytoplasm. Further research will be required to ascertain whether the Lys154Ala mutation underlies the defective closing of the external gate, causing the observed increase in external Km.

Collectively, our studies provide compelling evidence that the side chain of Lys154 is crucial for maintaining the stability of TM1, enabling its inherent dynamic gating between open and closed gate states, and facilitating that substrate binding biases the conformational equilibrium towards the closed gate structure (Fig 6). Akin to the role of Na2 in NSS transporters (Malinauskaite *et al*, 2014; Stolzenberg *et al*, 2017), the closed conformation sampling is facilitated by the hydrogen bonding between the conserved Lys154 in TM5 and Gly15 in TM1a. When this interaction is disrupted, as seen in the Lys154Ala mutation, it impedes the natural tilting of TM1a and the closing induced by the substrate at the cytoplasmic gate.

**Figure 6.**
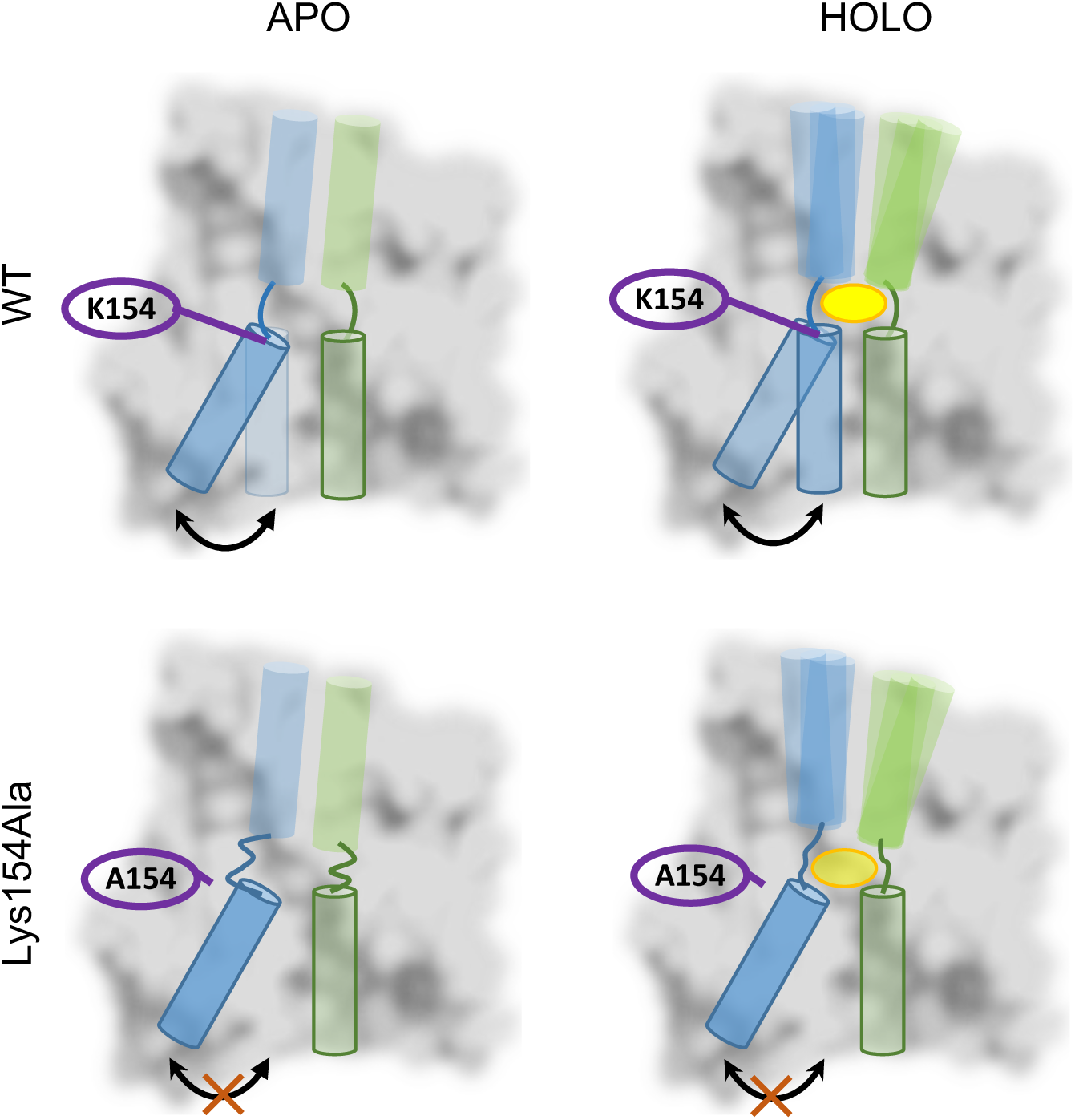
Schematic representation illustrating the movement of TM1a und apo (left) and substrate-bound (right) conditions, emphasizing the crucial role of Lys154. The diagram highlights the pivotal function of Lys154 (K154) in facilitating this process. Arrows indicate the transitions between different conformational states, reflecting the asymmetric equilibrium identified via single-molecule Förster resonance energy transfer (smFRET) and molecular dynamics (MD) studies. This equilibrium corresponds to distinct intracellular and extracellular K_m_ values and a significant reduction in V_max_, as previously documented (Bartoccioni et al., 2019; Errasti-Murugarren et al., 2019). The illustration provides a conceptual overview of the conformational changes facilitated by Lys154, underscoring their role in substrate binding and transport in BasC.

## Materials and Methods

### Mutagenesis

Point mutations in BasC were performed through QuickChange site-directed mutagenesis and verified by Sanger sequencing. TM1a-TM4 double cysteine variants were generated on a Cys-less variant: Ile7Cys-Thr120Cys-Cys427Ala.

### BasC expression and purification

A more detailed protocol of BasC purification was published in (Bartoccioni *et al*, 2019). Briefly, membranes of *E. coli* induced ON at 37 °C with BasC-3C-GFP-His plasmid were thawed and diluted with TB (20 mM Tris, 150 mM NaCl) until 3 mg/ml and incubated for 1 h at 4 °C with detergent (1 % DDM for proteoliposome (PL) preparation and smFRET assays or 2 % DM for structural analysis -Xray/Cryo-EM). Non-solubilized membranes were discarded by ultracentrifugation (200,000 x g, 1 h, 4 °C). Solubilized BasC-GFP was bound to Ni-NTA Agarose (Qiagen) previously equilibrated with DM or DDM buffer (TB with 0.06 % DDM or 0.17 % DM, respectively). Resin was first washed 3 x 10 CV with DM or DDM buffer with increasing concentrations of imidazole (10, 20 and 30 mM, respectively). Next, protein was eluted with DM or DDM buffer with 300 mM imidazole for PLs preparation. On the other hand, 3C protease elution was performed on column for structural analysis and smFRET assays. Here, resin was equilibrated with 10 CV of 3C buffer (DDM buffer supplemented with 1 mM DTT and 1 mM EDTA). His-tagged HRV-3C protease was added to the resin and incubated for 3 h (for FRET experiments) or overnight (for structural studies) at 4 °C.

### Nanobodies (Nbs) generation, expression and purification

Nbs were generated as described previously (Pardon *et al*, 2014; Errasti-Murugarren *et al*, 2019). One llama was immunized through six weekly injections of purified BasC reconstituted in E. coli polar lipid proteoliposomes. A panning experiment on BasC resulted in 29 specific Nb families. Nb53, Nb58, Nb71, and Nb74 were selected based on their transport-blocking activity as explained before. All used Nbs were expressed and purified in pMESy4 as inducible periplasmic proteins in E. coli WK6 strain. They produced milligram amounts and were purified to >95% using immobilized Ni-NTA chromatography (Qiagen) from a periplasmic extract of a 1 L culture. Purified Nbs (5–10 mg/ml) in 20 mM Tris-Base, NaCl 150 mM, pH 7.4, were frozen in liquid nitrogen and stored at −80°C before use. For structural purposes, we purified BasC complexed with Nbs as in Errasti-Murugarren *et al*. (2019). Ni-NTA pure BasC and desired Nbs were incubated in a molar ratio of 1:1.4 (BasC:Nb) for 1 h on ice. The complex was separated from Nb excess by size exclusion chromatography (SEC) on a Superdex-200 10/300 column (GE Healthcare). To concentrate we used protein concentrators with a cut-off of 100 kDa (Amicon) to avoid detergent concentration.

### Labelling

On-column labelling started with the same purification protocol as described previously but with the addition of 10 mM DTT from the lysis to labelling step. Before 3C elution, a 10 CV wash with DDM buffer (without DTT) was performed. Immediately after, maleimide fluorophores (Alexa Fluor 546 and Alexa Fluor 647 or the pair sulfo-Cy3 and sulfo-Cy5) were added in a BasC:dye molar ratio of 1:5. Incubation was carried out at 4 °C overnight in rotation and under light protection. The resin was washed three times using 10 CV with DDM buffer to remove free dyes, and 3C protease elution was done as described before. Labelled protein was finally separated from remaining free dye by SEC in a Superdex 200 10/300 GL column using DDM buffer and measuring absorbance at 554, 647, 561 and 650 nm for sulfo-Cy3, sulfo-Cy5, Alexa 546 and Alexa 647 respectively.

### ALEX-smFRET microscopy

A full description of the in-house confocal microscope and data analysis procedures for smFRET was provided previously and was adapted closely to (Gebhardt *et al*, 2021). Briefly, all samples were examined by focusing the excitation/observation volume of a confocal microscope into a ∼100 μl sample drop containing ∼50 pM of BasC in DDM buffer on a glass coverslip. Before data recording, the coverslips were coated for >60 s with 1 mg/mL bovine serum albumin to prevent fluorophore and/or protein adsorption.

Each experimental condition (apo, ligand etc.) was then studied for between 30 and 120 min, depending on the fraction of donor- and acceptor-containing BasC molecules in relation to donor- and acceptor-only ones. We also performed DNA controls with the same fluorophore pairs as for the proteins in DDM buffer to verify proper microscope alignment and to exclude buffer interferences (Appendix Fig S1).

For alternating laser excitation (Kapanidis *et al*, 2004; Hohlbein *et al*, 2014) we used two laser sources for sample excitation with electronic modulation: a 532 nm diode laser at 60 μW (OBIS 532-100-LS, Coherent, USA) and a diode laser at 640 nm (OBIS 640- 100-LX, Coherent, USA) at 25 or 30 μW for Alexa 546-Alexa 647 and sCy3-sCy5 dye-pairs, respectively. The laser light was guided into an epi-illuminated confocal microscope body (Olympus IX71, Hamburg, Germany) by a dual-edge beamsplitter (ZT532/640rpc, Chroma/AHF, Germany) and focused to a diffraction-limited excitation spot by a water immersion objective (UPlanSApo 60x/1.2w, Olympus Hamburg, Germany). The emitted fluorescence was collected through the same objective, spatially filtered using a pinhole with a diameter of 50 μm and spectrally split into donor and acceptor channel by a single-edge dichroic mirror H643 LPXR (AHF). Fluorescence emission was filtered (donor: BrightLine HC 582/75 (Semrock/AHF), acceptor: Longpass 647 LP Edge Basic (Semroch/AHF)) and focused onto avalanche photodiodes (SPCM-AQRH-64, Excelitas). The detector outputs were recorded by a NI-Card (PCI-6602, National Instruments, USA).

Data analysis and plottings was performed using an in-house written software package as described in (de Boer *et al*, 2019; Gouridis *et al*, 2015). Single-molecule events were identified using an all-photon-burst-search APBS, (Eggeling *et al*, 1998) with a threshold of 15, a time window of 500 μs and a minimum total photon number of 50, with an additional per-bin with a minimum total photon number between 100 and 150 for inclusion of the burst in the histogram. Comparisons between absolute results and mean FRET efficiencies E*, which are setup- and alignment-dependent, were made only when data were collected on the same day.

### Influx/Efflux assays in BasC Proteoliposomes

For transport measurements, PLs were prepared as previously described (Bartoccioni *et al*, 2019). Imidazole-eluted BasC-GFP protein was reconstituted in *E. coli* polar lipid extract (Sigma-Aldrich). Lipids were dried under nitrogen flow and resuspended in Tris buffer by bath sonication. Purified BasC protein was added to reach the desired protein to lipid ratio of 1:100 (w:w). To destabilize the liposomes, 1.25 % ß-D-octylglucoside was added and the mixture was incubated on ice for 30 min. PLs were formed by dialysis for 24-48 h at 4 °C against 100 volumes of Tris buffer. PL suspensions were aliquoted, frozen in liquid nitrogen, and stored at −80 °C until use.

L-alanine (A7627, Sigma-Aldrich), L-serine (S4500, Sigma-Aldrich), or L-glutamine (G8540, Sigma-Aldrich), along with 0.05 µCi/µl of L-[3H]-serine (Perkin Elmer) only for efflux experiments, were added to the PL suspension at the desired concentration. The mixture underwent six freeze/thaw cycles in liquid nitrogen to load the PLs. The excess liposomal solution was removed through ultracentrifugation (200,000 × g for 1 h at 4 °C), and the PLs were subsequently resuspended to one-third of the initial volume using Tris buffer. When amino acid concentrations exceeded 10 mM, either inside or outside the PLs, an equal concentration of D-mannitol (M4125, Sigma Aldrich) was used on the opposite side of the PLs to maintain osmotic balance.

Influx and efflux assays were initiated after mixing 20 μl of PLs solution with 180 μl of influx transport buffer (10 μM L-serine in Tris buffer with and 0.5 µCi of L-[3H]-serine (Perkin Elmer) or 4 mM cold amino acid for efflux experiments. Transport was allowed at RT and stopped in 5 s, adding 3 ml of ice-cold Tris buffer and by filtrating the PLs through 0.45 μm pore-size membrane filters (Sartorius Stedim Biotech). Filters were washed twice with 3 ml of Tris buffer and allowed to dry. Trapped radioactivity was then counted. Transport measured in PLs containing no amino acid was subtracted from each data point to calculate the net exchange. Transport measurements were normalized to the BasC protein concentration by SDS-PAGE.

### X-ray crystallography

Protein drops from purified BasC-Nb53-Nb58 complex were settled in a sitting-drop 96 wells plate using Oryx8 protein dispenser (Douglas Instruments). Crystallization was performed by vapour diffusion at 4 °C by mixing equal volumes of protein and reservoir solution containing 27–29 % PEG 400 and 0.1 M ammonium acetate pH 8 (0.1 M glycine pH 9 for HiLiDe crystals). Crystals typically appeared after 3 days, reaching a maximum size after 7-10 days. Crystals were cryoprotected by soaking in 33 % PEG 400 and then flash-cooled in liquid nitrogen and diffracted at the ALBA synchrotron light source (Cerdanyola del Vallès, Spain) and at the European Synchrotron Radiation Facility (Grenoble, France).

### X-ray data processing

Data were processed with AutoProc toolbox (Vonrhein *et al*, 2011). Phases were obtained by molecular replacement using PHASER (McCoy, 2007) and the structures of BasC and Nb74 (PDB ID: 6F2G) as templates. Model building into the electron density map was performed in COOT (Emsley *et al*, 2010), with a few cycles of structure refinement carried out in REFMAC5 (Murshudov *et al*, 2011). Images were prepared using Open-Source PyMol (The PyMOL Molecular Graphics System, Version 2.0 Schrödinger, LLC.) (DeLano, 2002).

### Cryo-EM

The specimen was vitrified using 3 μL of freshly purified BasC-Nb71 complex in 0.17% DM applied to glow-discharged Quantifoil R 0.6/1 Cu 300 mesh grids (Electron Microscopy Sciences), at a concentration of 2.2 mg/ml. Grids were blotted for 2 s under 95% humidity and plunge frozen in liquid ethane using a Vitrobot Mark IV (Thermo Fisher Scientific). Cryo-EM datasets were collected on a Titan Krios electron microscope operating at 300 kV and equipped with Thermo Fisher Falcon 4 direct electron detector at the facilities of the University of Leeds. A total of 6,756 movies were collected in .eer format. The set-up used a 130k magnification, which corresponded to 0.91 Å/pix. The defocus applied covered from -3.3 um to -1.8 μm, and the total electron dose was 34.83 e-/Å^2^.

### Cryo-EM image processing

Movies were imported to RELION4 (Kimanius *et al*, 2021) and aligned using MotionCor2 (Zheng *et al*, 2017) with 35 patches per image and 31 frames per group, applying dose weighting. Contrast transfer function (CTF) parameters were determined using the CTFFIND-4.1 approach (Rohou & Grigorieff, 2015). Particles were selected using Topaz, after training the neural network using a∼1,000 manually selected particles in a subset of 100 randomly selected micrographs (Bepler *et al*, 2020). Subsequent image processing was performed using RELION4, and cryoSPARC (Punjani *et al*, 2017). Initially, 590,169 particles were extracted and subjected to several rounds of reference-free 2D classification, obtaining 409,984 particles. Selected particles were used to generate 3D reconstructions, leading to a consensus volume of >160,000 particles where all TMs were already visible. To recover good particles, iterative rounds of heterogeneous refinement were performed using cryoSPARC. Thus, an ab initio reconstruction was used to clean the particles in 3D, selecting a final subset of 169,672 particles. By non-homogeneous refinement (Punjani *et al*, 2020), the density from the micelle was removed and a map of ∼6.0 Å resolution was generated. Volumes were visualized in UCSF Chimera (Pettersen *et al*, 2004), which was also used to fit the available BasC (PDB ID 6F2G) into the density.

### Molecular dynamics simulations and post-processing analysis

All the simulated systems presented here were prepared starting from the BasC crystal structure (PDB ID: 6F2W) (Errasti-Murugarren *et al*, 2019) in the open-inward conformation. By either mutating Lys154Ala or removing the substrate (2AIB), we obtained 4 distinct BasC systems: 1) WT in apo condition; 2) WT in holo condition; 3) Lys154Ala mutated in apo condition; and 4)Lys154Ala mutated in holo condition.

All these systems were inserted in 150 x 150 Å^2^ lipid membrane perpendicular to z-axis whose concentration mimics the *E. coli* polar lipid extract (Sigma-Aldrich) used in PLs transport assay (70 % POPG, 10 % Cardiolipin (TMCL) and 20% POPE.). Above and below the lipid bilayer, the simulation boxes were filled with TIP3P water molecules and a few Na+ and K-ions to neutralize the systems and bring the total ion concentration to 0.15 uM. The resulting heights along the z-axis were 100 Å. We used the CHARMM-GUI web-server to set up the starting configurations (Jo *et al*, 2008). The simulations were performed using GROMACS 2020.2 code (Abraham *et al*, 2015) and CHARMM 36 forcefield (Best *et al*, 2012). LINCS (Hess *et al*, 1997) algorithm was used to constrain all the bonds involving hydrogen atoms, thereby allowing us to employ a 2 fs step to integrate the Newton equations. The Mesh Ewald method was used to account for long-range interactions with a real-space cut-off of 12 Å. The first minimization employing a steepest descendent algorithm (Steinmetz, 1966) was followed by an equilibration step consisting of an NPT ensemble 100 ns run with a semi-isotropic Berendsen barostat (Berendsen *et al*, 1984) to maintain the pressure at 1 atm with a coupling constant of 0.6 ps. The temperature, set at 310 K, was controlled using the Velocity rescaling algorithm with a 0.4 ps coupling constant (Bou-Rabee, 2014). The production step was a 500 ns long NPT ensemble simulation using a semi-isotropic Parrinello-Rahman barostat (Parrinello & Rahman, 1981) with a 0.6 ps coupling constant and a Nose-Hoover thermostat (Evans & Holian, 1985) with a 0.4 ps coupling constant to fix pressure and temperature at 1 atm and 310K, respectively. We performed three separate replicas for each system. All the analyses presented in this work were carried out using either VMD (Humphrey *et al*, 1996) software or in-house scripts.

## Data Availability

- Cryo-EM volume (Fig 3C): deposit in process in EMDB
- X-ray low resolution data (Fig 3D) (Provisional link) https://data.mendeley.com/preview/4hyrf9x778?a=12dd5e92-63ba-484a-b983-70517254f3ca
- Scripts for generating MD histograms: Github (https://github.com/LucaMaggi19/Histogram-)

## Acknowledgements

This work was funded by the Spanish Ministry of Science and Innovation (PID2021-122802OB-I00) and the Catalan Government (grant 2021 SGR 01281). We thank INSTRUCT-ERIC for providing Nanobodies (PID1176) and access to 300kV Titan Crios of Leeds, UK (PID17338). We acknowledge the European Synchrotron Radiation Facility (ESRF) for provision of synchrotron radiation facilities, and we would like to thank Julien Orlans for assistance and support in using beamline ID30A3. We thank Nick Berrow (Protein Expression Core Facility; IRB Barcelona) for providing us with HRV-3C protease, and Joan Pous from the IRB Barcelona/IBMB-CSIC Crystallization Platform. We also thank Jasminka Boskovic and Johanne LeCoq from the Electron Microscopy Unit at CNIO for preparing and visualizing the grids and sending samples to Leeds. We gratefully acknowledge institutional funding from the Spanish State Research Agency of the Spanish Ministry of Science and Innovation—Programa Estatal de Fomento de la Investigación Científica y Técnica de Excelencia—Centres of Excellence “Severo Ochoa” CEX2019-000891-S and CEX2019-000913-S. O.L. laboratory also had the support from the National Institute of Health Carlos III to CNIO. IRB Barcelona is a member of the Centres de Recerca de Catalunya (CERCA) System of the Catalan Government. J.F. is supported by a Centro de Investigación Biomédica en Red de Enfermedades Raras (CIBERER) contract and A.N. was supported by FI-AGAUR Fellowship. M. M. M. is supported by La Caixa (LCF/PR/HR20/52400017 to O.L. and M.P.). Research in the laboratory of T.C. was supported by the European Comission (ERC-STG 638536 – SM-IMPORT), Deutsche Forschungsgemeinschaft (GRK2062, project C03 to T.C.; SFB863, project A13 to T.C. and the Center for Nanosicence (CeNS). L.M. and M.O. acknowledge support by: Spanish Ministerio de Ciencia e Innovación [PID2020-116620GB-I00, RTI2018-096704-B-100, PID2021-122478NB-I00]; Center of Excellence for HPC H2020 European Commission; BioExcel-3. Centre of Excellence for Computational Biomolecular Research [823830]; Catalan SGR and the Instituto de Salud Carlos III Instituto Nacional de Bioinformatica [ISCIII PT 17/0009/0007 co-funded by the Fondo Europeo de Desarrollo Regional]; European Regional Development Fund under the framework of the ERFD Operative Programme for Catalunya, the Catalan Government AGAUR [SGR2017-134]; European Union MDDB: Molecular Dynamics Data Bank. The European Repository for Biosimulation Data [101094651]. The project used computer resources from the Barcelona Supercomputing Centre (BSC). We thank Gabriel G. Moya-Munoz, Atieh Aminian Jazi and Felix Pioch for support of this study. Finally, we are grateful to Pol Torrents, Laia Bekius and Javier Fernández for help during their student internships.

## Expanded View Figure legends

**Figure EV1.**
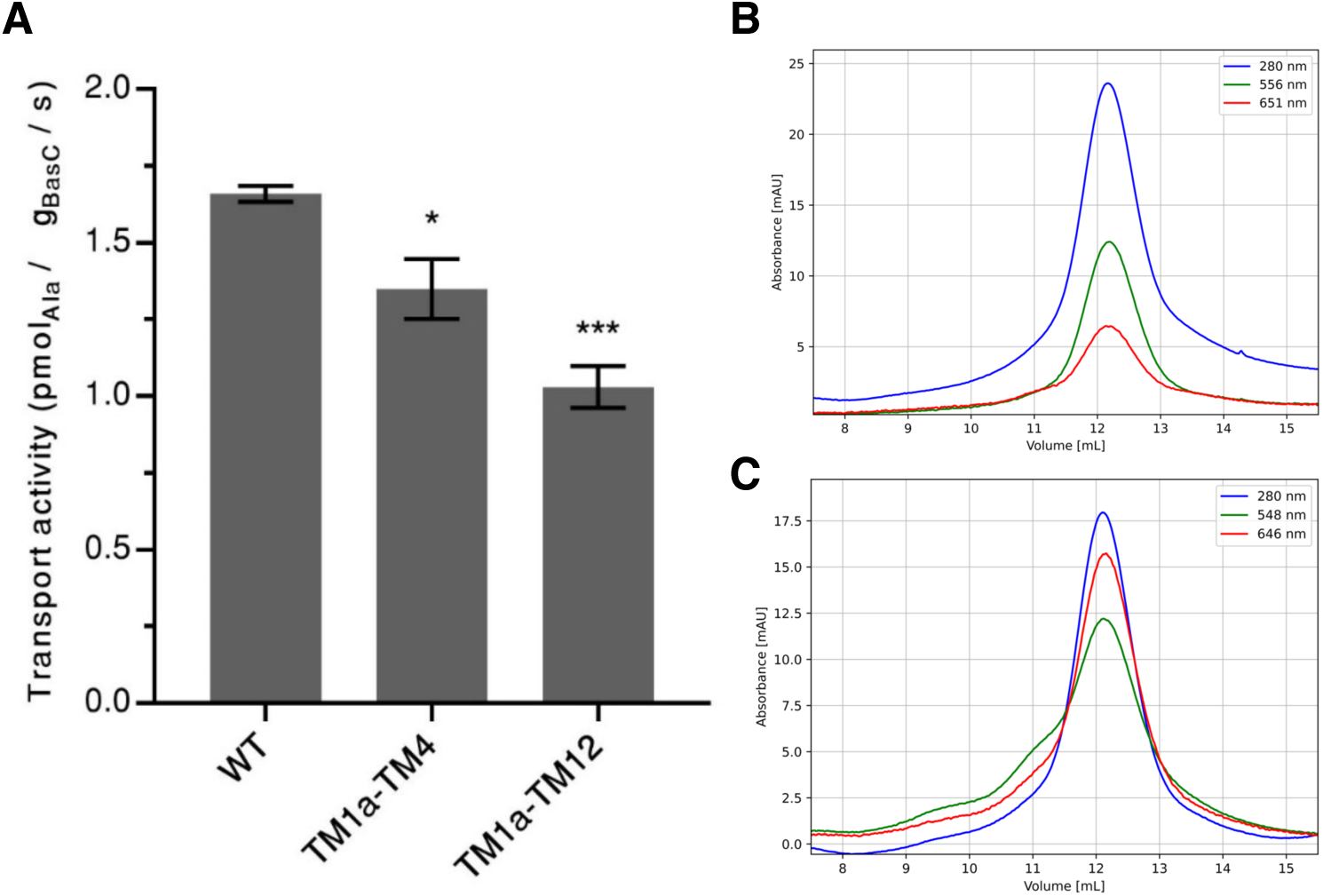
Biochemical characterization of BasC cysteine variants. **A.** Relative influx transport activity of BasC double-Cys variants. 5-seconds influx of 10 μM L-[3H]-Ser (1 μCi/μl) into BasC-GFP-PLs (BasC-Green Fluorescent Protein PLs) containing 4 mM L-alanine was measured for BasC WT and double-Cys variants TM1a-4 and TM1a-12 reconstituted in PLs. Data represents mean ± standard error of the mean (SEM) from triplicates of a single experiment. T-tests were performed comparing transport activity data of double-Cys mutants with WT (*p < 0.05, ***p < 0.005). **B.** and **C**. SEC elution profiles of TM1a-TM12 (B) and TM1a-TM4 (C) BasC variants labelled with Alexa 546/Alexa 647 (B) and sulfo-Cy3/sulfo-Cy5 (C), respectively. The protein is detected by 280 nm absorbance (blue), while donor and acceptor dyes are seen at 556 and 651 nm for Alexa 546/Alexa 647 (red and green) or 548 and 646 nm (red and green) for sulfo-Cy3/sulfo-Cy5.

**Figure EV2.**
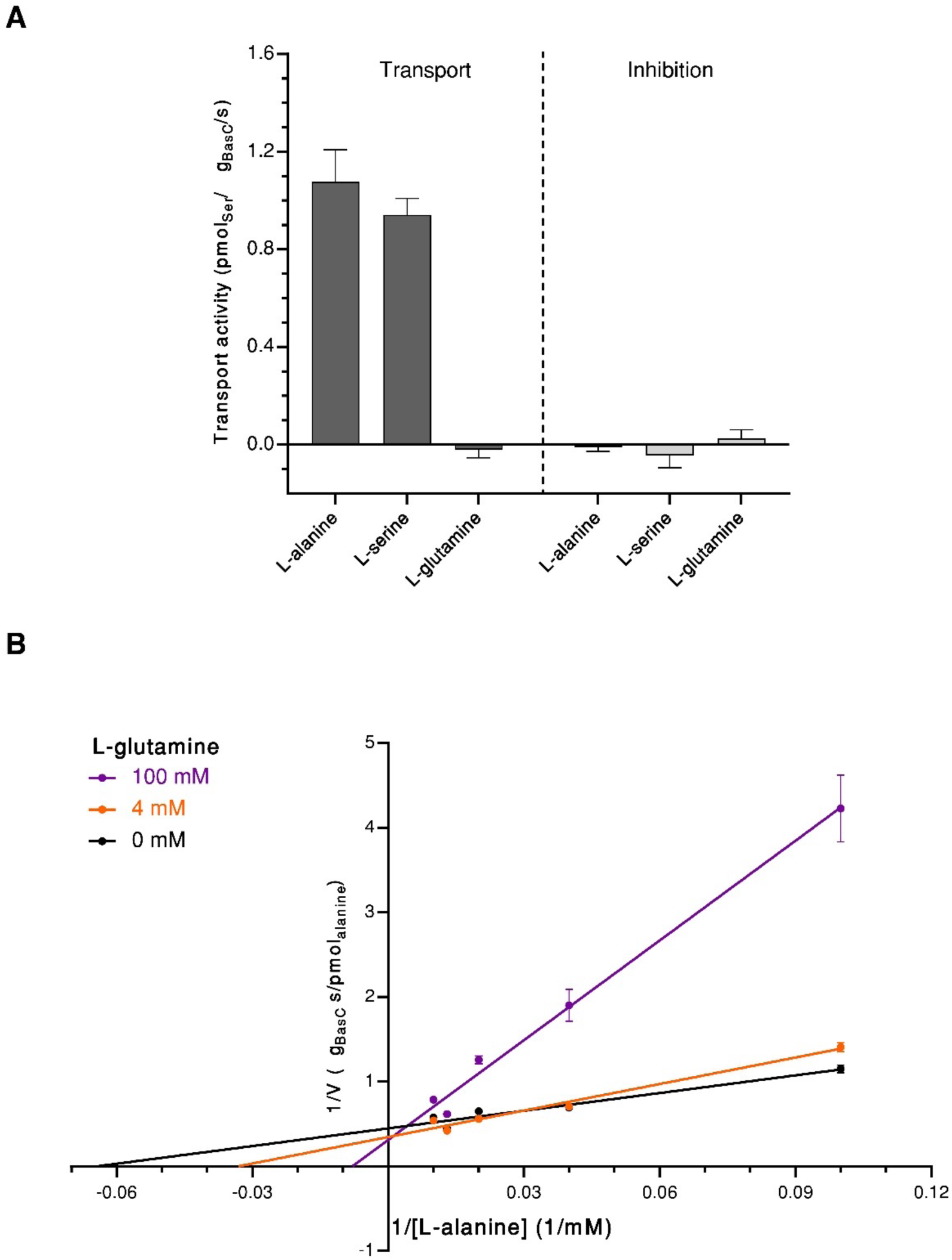
BasC reconstituted in liposomes transports L-alanine and L-serine but not the competitive inhibitor L-glutamine. **A.** Substrate specificity at high concentrations of amino acids: BasC influx transport of 10 μM L-[3H]-Ser (1 μCi/μl) into BasC-GFP-PLs (BasC-Green Fluorescent Protein PLs) containing 100 mM of the indicated amino acid (AA). Data represent mean ± standard error of the mean (SEM) from three independent experiments. **B.** The Lineweaver-Burk plot illustrates competitive inhibition of BasC by L-glutamine. Increasing concentrations of L-glutamine caused to an increase in K_m_ for L-alanine while the V_max_ was unaffected. L-alanine was titrated in the absence and presence of L-Gln at 4 and 100 mM. Data represent mean ± SEM from triplicates of a single experiment. Similar results were obtained in a second experiment with different concentrations.

**Figure EV3.**
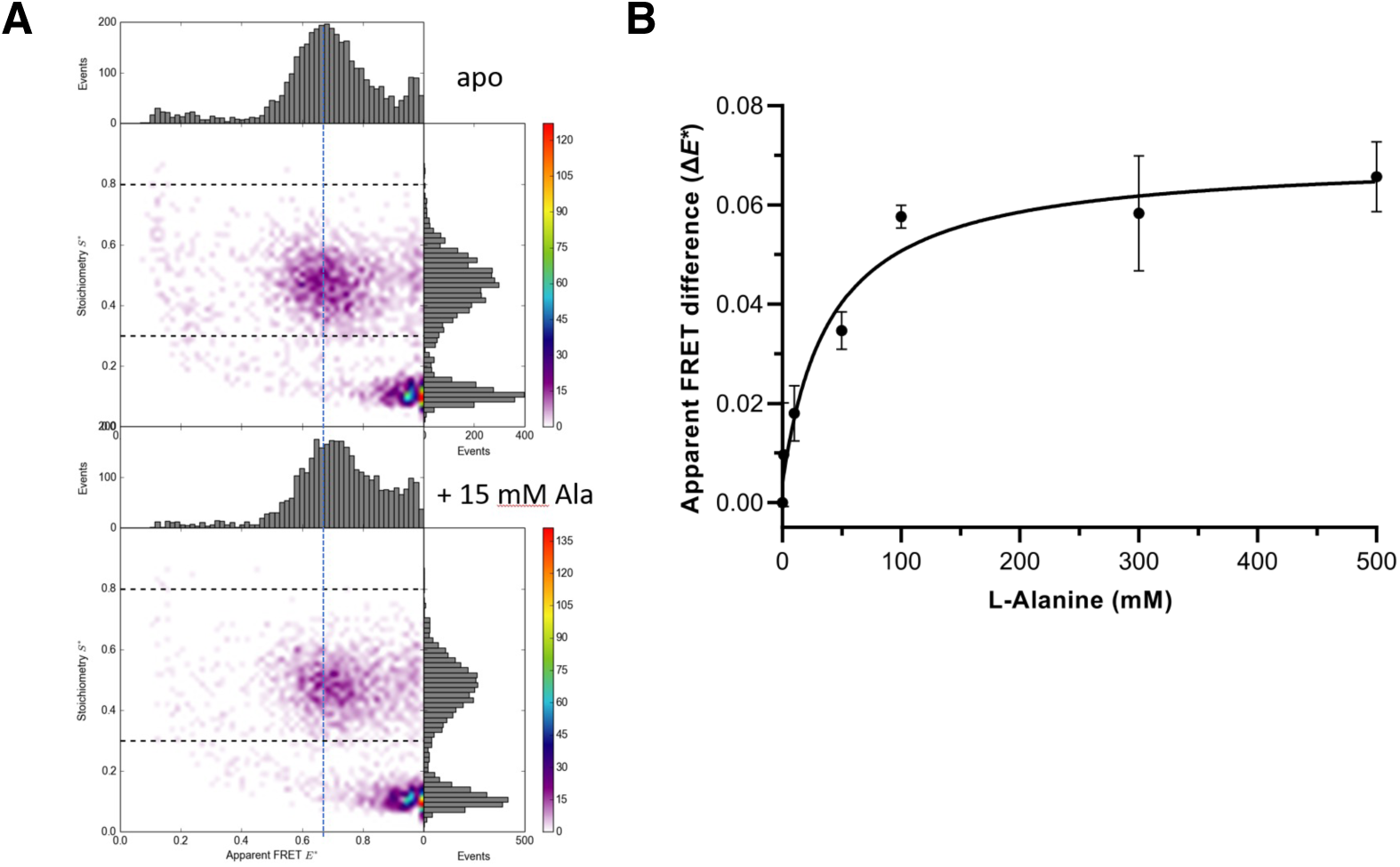
**A. TM1a-TM12a labeled with sCy3 and sCy5 showed a similar behavior to TM1a-TM4**, with a peak exhibiting a tendency to shift molecules to higher E* FRET when substrate was added. However, we were not able to observe two populations as for the Alexa dyes. **B. L-Alanine effects in smFRET experiments.** Effect of L-Alanine concentrations on smFRET data for BasC double-cysteine variant TM1a-4 labelled with sCy3 and sCy5 in DDM. Apparent FRET differences between apo and holo conditions upon increasing L-Ala concentrations. Mean ± SEM values of 1E* are presented from three independent experiments. Non-linear regression fitting was performed using GraphPadPrism 9.5.1, and EC50 for the smFRET effect was calculated to be 40 ± 13 mM.

**Figure EV4.**
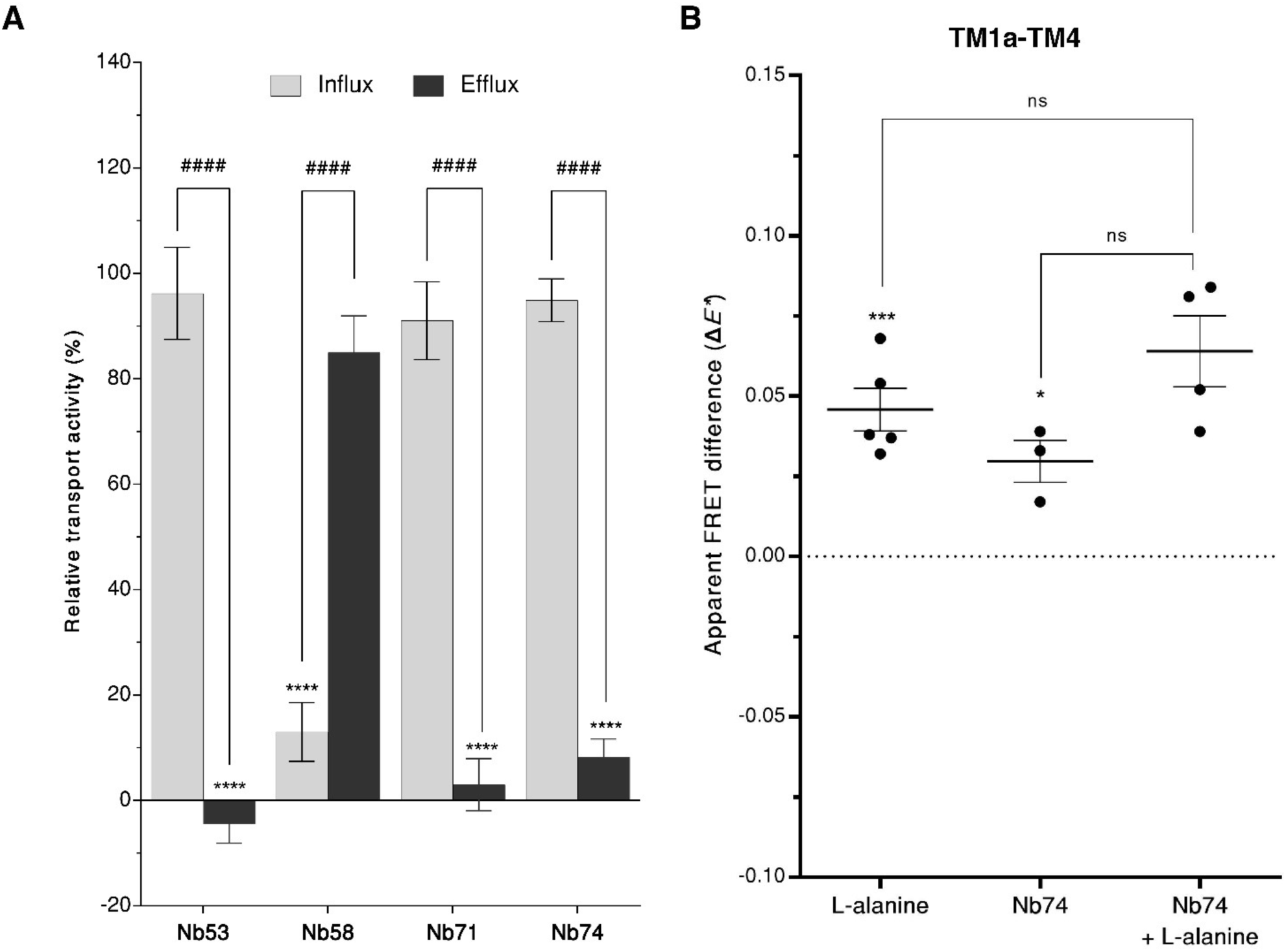
Quantification of Nb74 effects on transport and smFRET. **A.** Relative transport influx and efflux activity screening for each BasC-Nb complex. Influx and efflux quantification was performed after 5 and 30 seconds, respectively on BasC-GFP reconstituted into PLs treated with Nb at concentration according to their effective inhibition concentrations, which was between 5 and 100 µM. Mean ± SEM are plotted for each Nb treatment for a total of minimum of three independent experiments with the exception of the Nb that did not show influx inhibition below 70% during the first two trials. Multiple Dunett test comparisons were performed for each Nb (influx or efflux) against relative activity of the non-treated sample. Multiple T-tests were performed between influx and efflux inhibition values within the same Nb. Statistical significance against the non-treated (100 %) is shown in asterisks and within the same Nb in hashes (**** or #### p.value < 0.0001). **B.** Differences in apparent smFRET Δ*E\** effects of the TM1a-TM4 pair with or without a pre-incubation with Nb74. Statistical analysis after incubation with the substrate was performed relative to apo measurement of the same day (ΔE* = 0, dashed horizontal line) with the absolute values. Mean ± SEM values of ΔE*are presented. P-paired T-tests were performed with the absolute *E\** values between apo measurements on the same day. P-values are indicated by asterisks (*** < 0.005, * < 0.05; and ns, non-statistically difference. Values for Nb74 or Nb74 plus L-alanine treatments showed a p-value of 0.06.

**Figure EV5.**
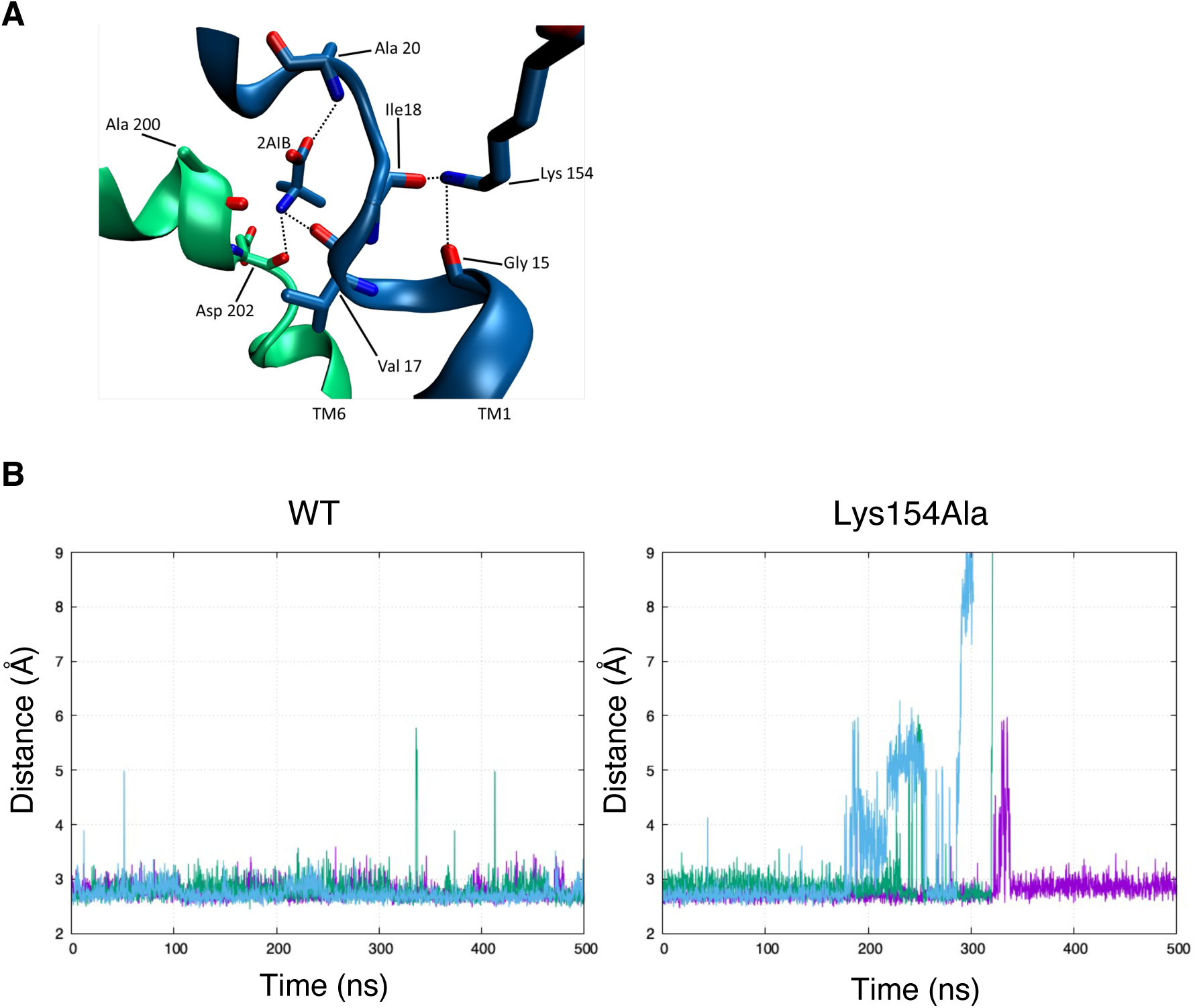
Molecular dynamics simulations and results of the BasC holo state. **A.** Representative snapshot from molecular dynamics simulations of the binding site of BasC wild type in the holo state. TM1 and TM6 are represented in blue and green, respectively. The dashed lines indicate hydrogen bonds with substrate, illustrating the spatial arrangement in the binding site and offering a detailed view of the molecular interactions. **B, C**. Distance between the Val 17 carbonyl oxygen (in blue) and the substrate (2AIB) nitrogen in three 500 ns replicas of (B) BasC wild type and (C) BasC Lys154Ala.

## References

Abraham MJ, Murtola T, Schulz R, Páll S, Smith JC, Hess B & Lindahl E (2015) GROMACS: High performance molecular simulations through multi-level parallelism from laptops to supercomputers. SoftwareX 1: 19–25

Agam G, Gebhardt C, Popara M, Mächtel R, Folz J, Ambrose B, Chamachi N, Chung SY, Craggs TD, de Boer M, et al (2023) Reliability and accuracy of single-molecule FRET studies for characterization of structural dynamics and distances in proteins. Nat Methods 20: 523–535

del Alamo D, Meiler J & Mchaourab HS (2022) Principles of Alternating Access in LeuT-fold Transporters: Commonalities and Divergences. Journal of Molecular Biology 434: 167746

Bartels K, Lasitza-Male T, Hofmann H & Löw C (2021) Single-Molecule FRET of Membrane Transport Proteins. ChemBioChem 22: 2657–2671

Bartoccioni P, Fort J, Zorzano A, Errasti-Murugarren E & Palacín M (2019) Functional characterization of the alanine-serine-cysteine exchanger of Carnobacterium sp AT7. Journal of General Physiology 151: 505–517

Bepler T, Kelley K, Noble AJ & Berger B (2020) Topaz-Denoise: general deep denoising models for cryoEM and cryoET. Nat Commun 11: 5208

Berendsen HJC, Postma JPM, van Gunsteren WF, DiNola A & Haak JR (1984) Molecular dynamics with coupling to an external bath. The Journal of Chemical Physics 81: 3684–3690

Best RB, Zhu X, Shim J, Lopes PEM, Mittal J, Feig M & MacKerell ADJr (2012) Optimization of the Additive CHARMM All-Atom Protein Force Field Targeting Improved Sampling of the Backbone ϕ, ψ and Side-Chain χ1 and χ2 Dihedral Angles. J Chem Theory Comput 8: 3257–3273

de Boer M, Gouridis G, Vietrov R, Begg SL, Schuurman-Wolters GK, Husada F, Eleftheriadis N, Poolman B, McDevitt CA & Cordes T (2019) Conformational and dynamic plasticity in substrate-binding proteins underlies selective transport in ABC importers. eLife 8: e44652

Bou-Rabee N (2014) Time Integrators for Molecular Dynamics. Entropy 16: 138–162

Debruycker V, Hutchin A, Masureel M, Ficici E, Martens C, Legrand P, Stein RA, Mchaourab HS, Faraldo-Gómez JD, Remaut H, et al (2020) An embedded lipid in the multidrug transporter LmrP suggests a mechanism for polyspecificity. Nat Struct Mol Biol 27: 829–835

DeLano WL (2002) Pymol: An open-source molecular graphics tool. CCP4 Newsletter On Protein Crystallography: 82–92

Drehmann P, Milanos S, Schaefer N, Kasaragod VB, Herterich S, Holzbach-Eberle U, Harvey RJ & Villmann C (2023) Dual Role of Dysfunctional Asc-1 Transporter in Distinct Human Pathologies, Human Startle Disease, and Developmental Delay. eNeuro 10

Dyla M, Terry DS, Kjaergaard M, Sørensen TL-M, Lauwring Andersen J, Andersen JP, Rohde Knudsen C, Altman RB, Nissen P & Blanchard SC (2017) Dynamics of P-type ATPase transport revealed by single-molecule FRET. Nature 551: 346–351

Eggeling C, Fries JR, Brand L, Günther R & Seidel CA (1998) Monitoring conformational dynamics of a single molecule by selective fluorescence spectroscopy. Proc Natl Acad Sci U S A 95: 1556–1561

Emsley P, Lohkamp B, Scott WG & Cowtan K (2010) Features and development of Coot. Acta Crystallogr D Biol Crystallogr 66: 486–501

Errasti-Murugarren E, Fort J, Bartoccioni P, Díaz L, Pardon E, Carpena X, Espino-Guarch M, Zorzano A, Ziegler C, Steyaert J, et al (2019) L amino acid transporter structure and molecular bases for the asymmetry of substrate interaction. Nat Commun 10: 1807

Espino Guarch M, Font-Llitjós M, Murillo-Cuesta S, Errasti-Murugarren E, Celaya AM, Girotto G, Vuckovic D, Mezzavilla M, Vilches C, Bodoy S, et al (2018) Mutations in L-type amino acid transporter-2 support SLC7A8 as a novel gene involved in age-related hearing loss. eLife 7: e31511

Evans DJ & Holian BL (1985) The Nose–Hoover thermostat. The Journal of Chemical Physics 83: 4069–4074

Feliubadaló L, Bisceglia L, Font M, Dello Strologo L, Beccia E, Arslan-Kirchner M, Steinmann B, Zelante L, Estivill X, Zorzano A, et al (1999) Recombinant families locate the gene for non-type I cystinuria between markers C13 and D19S587 on chromosome 19q13 1. Genomics 60: 362–365

Fotiadis D, Kanai Y & Palacín M (2013) The SLC3 and SLC7 families of amino acid transporters. Molecular Aspects of Medicine 34: 139–158

Gauthier-Coles G, Fairweather SJ, Bröer A & Bröer S (2023) Do Amino Acid Antiporters Have Asymmetric Substrate Specificity? Biomolecules 13: 301

Gauthier-Coles G, Vennitti J, Zhang Z, Comb WC, Xing S, Javed K, Bröer A & Bröer S (2021) Quantitative modelling of amino acid transport and homeostasis in mammalian cells. Nat Commun 12: 5282

Gebhardt C, Lehmann M, Reif MM, Zacharias M, Gemmecker G & Cordes T (2021) Molecular and Spectroscopic Characterization of Green and Red Cyanine Fluorophores from the Alexa Fluor and AF Series. Chemphyschem 22: 1566–1583

Gouridis G, Schuurman-Wolters GK, Ploetz E, Husada F, Vietrov R, de Boer M, Cordes T & Poolman B (2015) Conformational dynamics in substrate-binding domains influences transport in the ABC importer GlnPQ. Nat Struct Mol Biol 22: 57–64

Ha T, Enderle T, Ogletree DF, Chemla DS, Selvin PR & Weiss S (1996) Probing the interaction between two single molecules: fluorescence resonance energy transfer between a single donor and a single acceptor. Proc Natl Acad Sci U S A 93: 6264–6268

Hellenkamp B, Schmid S, Doroshenko O, Opanasyuk O, Kühnemuth R, Rezaei Adariani S, Ambrose B, Aznauryan M, Barth A, Birkedal V, et al (2018) Precision and accuracy of single-molecule FRET measurements—a multi-laboratory benchmark study. Nat Methods 15: 669–676

Hess B, Bekker H, Berendsen HJC & Fraaije JGEM (1997) LINCS: A linear constraint solver for molecular simulations. Journal of Computational Chemistry 18: 1463–1472

Hohlbein J, Craggs TD & Cordes T (2014) Alternating-laser excitation: single-molecule FRET and beyond. Chem Soc Rev 43: 1156–1171

Humphrey W, Dalke A & Schulten K (1996) VMD: Visual molecular dynamics. Journal of Molecular Graphics 14: 33–38

Husada F, Bountra K, Tassis K, Boer M, Romano M, Rebuffat S, Beis K & Cordes T (2018) Conformational dynamics of the ABC transporter McjD seen by single-molecule FRET. EMBO J 37

Jeckelmann J-M, Lemmin T, Schlapschy M, Skerra A & Fotiadis D (2022) Structure of the human heterodimeric transporter 4F2hc-LAT2 in complex with Anticalin, an alternative binding protein for applications in single-particle cryo-EM. Sci Rep 12: 18269

Jo S, Kim T, Iyer VG & Im W (2008) CHARMM-GUI: A web-based graphical user interface for CHARMM. Journal of Computational Chemistry 29: 1859–1865

Jungnickel KEJ, Parker JL & Newstead S (2018) Structural basis for amino acid transport by the CAT family of SLC7 transporters. Nature Communications 9: 1–12

Kanai Y (2022) Amino acid transporter LAT1 (SLC7A5) as a molecular target for cancer diagnosis and therapeutics. Pharmacology & Therapeutics 230: 107964

Kapanidis AN, Lee NK, Laurence TA, Doose S, Margeat E & Weiss S (2004) Fluorescence-aided molecule sorting: analysis of structure and interactions by alternating-laser excitation of single molecules. Proc Natl Acad Sci U S A 101: 8936– 8941

Khan JA, Sohail A, Jayaraman K, Szöllősi D, Sandtner W, Sitte HH & Stockner T (2020) The Amino Terminus of LeuT Changes Conformation in an Environment Sensitive Manner. Neurochem Res 45: 1387–1398

Kimanius D, Dong L, Sharov G, Nakane T & Scheres SHW (2021) New tools for automated cryo-EM single-particle analysis in RELION-4.0. Biochem J 478: 4169–4185

Knöpfel EB, Vilches C, Camargo SMR, Errasti-Murugarren E, Stäubli A, Mayayo C, Munier FL, Miroshnikova N, Poncet N, Junza A, et al (2019) Dysfunctional LAT2 Amino Acid Transporter Is Associated With Cataract in Mouse and Humans. Front Physiol 10: 688

Koppula P, Zhuang L & Gan B (2021) Cystine transporter SLC7A11/xCT in cancer: ferroptosis, nutrient dependency, and cancer therapy. Protein & cell 12: 599–620

Krishnamurthy H, Piscitelli CL & Gouaux E (2009) Unlocking the molecular secrets of sodium-coupled transporters. Nature 459: 347–355

Lasitza-Male T, Bartels K, Jungwirth J, Wiggers F, Rosenblum G, Hofmann H & Löw C (2020) Membrane Chemistry Tunes the Structure of a Peptide Transporter. Angewandte Chemie International Edition 59: 19121–19128

Lee Y, Jin C, Ohgaki R, Xu M, Ogasawara S, Warshamanage R, Yamashita K, Murshudov G, Nureki O, Murata T, et al (2023) Structural basis of anticancer drug recognition and amino acid transport by LAT1. 2023.12.03.567112 doi:10.1101/2023.12.03.567112 [PREPRINT]

Lee Y, Wiriyasermkul P, Jin C, Quan L, Ohgaki R, Okuda S, Kusakizako T, Nishizawa T, Oda K, Ishitani R, et al (2019) Cryo-EM structure of the human L-type amino acid transporter 1 in complex with glycoprotein CD98hc. Nat Struct Mol Biol 26: 510–517

Lee Y, Wiriyasermkul P, Kongpracha P, Moriyama S, Mills DJ, Kühlbrandt W & Nagamori S (2022) Ca2+-mediated higher-order assembly of heterodimers in amino acid transport system b0,+ biogenesis and cystinuria. Nat Commun 13: 2708

Lerner E, Barth A, Hendrix J, Ambrose B, Birkedal V, Blanchard SC, Börner R, Sung Chung H, Cordes T, Craggs TD, et al (2021) FRET-based dynamic structural biology: Challenges, perspectives and an appeal for open-science practices. eLife 10: e60416

Lerner E, Cordes T, Ingargiol A, Alhadid Y, Chung S, Michalet X & Weiss S (2018) Toward dynamic structural biology: Two decades of single-molecule Förster resonance energy transfer. Science 359: eaan1133

Malinauskaite L, Quick M, Reinhard L, Lyons JA, Yano H, Javitch JA & Nissen P (2014) A mechanism for intracellular release of Na+ by neurotransmitter/sodium symporters. Nat Struct Mol Biol 21: 1006–1012

McCoy AJ (2007) Solving structures of protein complexes by molecular replacement with *Phaser*. Acta Crystallographica Section D Biological Crystallography 63: 32–41

Meier C, Ristic Z, Klauser S & Verrey F (2002) Activation of system L heterodimeric amino acid exchangers by intracellular substrates. EMBO J 21: 580–589

Murshudov GN, Skubák P, Lebedev AA, Pannu NS, Steiner RA, Nicholls RA, Winn MD, Long F & Vagin AA (2011) REFMAC 5 for the refinement of macromolecular crystal structures. Acta Crystallographica Section D Biological Crystallography 67: 355–367

Pardon E, Laeremans T, Triest S, Rasmussen SGF, Wohlkönig A, Ruf A, Muyldermans S, Hol WGJ, Kobilka BK & Steyaert J (2014) A general protocol for the generation of Nanobodies for structural biology. Nat Protoc 9: 674–693

Parker JL, Deme JC, Kolokouris D, Kuteyi G, Biggin PC, Lea SM & Newstead S (2021) Molecular basis for redox control by the human cystine/glutamate antiporter system xc−. Nat Commun 12: 1–11

Parrinello M & Rahman A (1981) Polymorphic transitions in single crystals: A new molecular dynamics method. Journal of Applied Physics 52: 7182–7190

Pettersen EF, Goddard TD, Huang CC, Couch GS, Greenblatt DM, Meng EC & Ferrin TE (2004) UCSF Chimera--a visualization system for exploratory research and analysis. J Comput Chem 25: 1605–1612

Punjani A, Rubinstein JL, Fleet DJ & Brubaker MA (2017) cryoSPARC: algorithms for rapid unsupervised cryo-EM structure determination. Nat Methods 14: 290–296

Punjani A, Zhang H & Fleet DJ (2020) Non-uniform refinement: adaptive regularization improves single-particle cryo-EM reconstruction. Nat Methods 17: 1214–1221

Rodriguez CF, Escudero-Bravo P, Díaz L, Bartoccioni P, García-Martín C, Gilabert JG, Boskovic J, Guallar V, Errasti-Murugarren E, Llorca O, et al (2021) Structural basis for substrate specificity of heteromeric transporters of neutral amino acids. Proc Natl Acad Sci U S A 118: e2113573118

Rohou A & Grigorieff N (2015) CTFFIND4: Fast and accurate defocus estimation from electron micrographs. J Struct Biol 192: 216–221

Rudnick G, Krämer R, Blakely RD, Murphy DL & Verrey F (2014) The SLC6 transporters: perspectives on structure, functions, regulation, and models for transporter dysfunction. Pflugers Arch 466: 25–42

Shi L, Quick M, Zhao Y, Weinstein H & Javitch JA (2008) The mechanism of a neurotransmitter:sodium symporter – inward release of Na+ and substrate is triggered by substrate in a second binding site. Mol Cell 30: 667–677

Sperandeo MP, Andria G & Sebastio G (2008) Lysinuric protein intolerance: update and extended mutation analysis of the SLC7A7 gene. Hum Mutat 29: 14–21

Steinmetz HL (1966) USING THE METHOD OF STEEPEST DESCENT. Ind Eng Chem 58: 33–39

Stolzenberg S, Li Z, Quick M, Malinauskaite L, Nissen P, Weinstein H, Javitch JA & Shi L (2017) The role of transmembrane segment 5 (TM5) in Na2 release and the conformational transition of neurotransmitter:sodium symporters toward the inward-open state. Journal of Biological Chemistry 292: 7372–7384

Tavoulari S, Margheritis E, Nagarajan A, DeWitt DC, Zhang Y-W, Rosado E, Ravera S, Rhoades E, Forrest LR & Rudnick G (2016) Two Na+ Sites Control Conformational Change in a Neurotransmitter Transporter Homolog. J Biol Chem 291: 1456–1471

Terry DS, Kolster RA, Quick M, LeVine MV, Khelashvili G, Zhou Z, Weinstein H, Javitch JA & Blanchard SC (2018) A partially-open inward-facing intermediate conformation of LeuT is associated with Na+ release and substrate transport. Nat Commun 9: 230

Torrents D, Mykkänen J, Pineda M, Feliubadaló L, Estévez R, Cid R de, Sanjurjo P, Zorzano A, Nunes V, Huoponen K, et al (1999) Identification of SLC7A7, encoding y+LAT-1, as the lysinuric protein intolerance gene. Nat Genet 21: 293–296

Vonrhein C, Flensburg C, Keller P, Sharff A, Smart O, Paciorek W, Womack T & Bricogne G (2011) Data processing and analysis with the autoPROC toolbox. Acta Crystallogr D Biol Crystallogr 67: 293–302

Wu D, Grund TN, Welsch S, Mills DJ, Michel M, Safarian S & Michel H (2020) Structural basis for amino acid exchange by a human heteromeric amino acid transporter. Proc Natl Acad Sci U S A 117: 21281–21287

Xiong G, Liu C, Yang G, Feng M, Xu J, Zhao F, You L, Zhou L, Zheng L, Hu Y, et al (2019) Long noncoding RNA GSTM3TV2 upregulates LAT2 and OLR1 by competitively sponging let-7 to promote gemcitabine resistance in pancreatic cancer. J Hematol Oncol 12: 97

Yamashita A, Singh SK, Kawate T, Jin Y & Gouaux E (2005) Crystal structure of a bacterial homologue of Na + /Cl - -dependent neurotransmitter transporters. Nature 437: 215–223

Yan R, Li Y, Müller J, Zhang Y, Singer S, Xia L, Zhong X, Gertsch J, Altmann K-H & Zhou Q (2021) Mechanism of substrate transport and inhibition of the human LAT1-4F2hc amino acid transporter. Cell Discov 7: 16

Yan R, Li Y, Shi Y, Zhou J, Lei J, Huang J & Zhou Q (2020a) Cryo-EM structure of the human heteromeric amino acid transporter b ^0,+^ AT-rBAT. Sci Adv 6: eaay6379

Yan R, Zhao X, Lei J & Zhou Q (2019) Structure of the human LAT1–4F2hc heteromeric amino acid transporter complex. Nature 568: 127–130

Yan R, Zhou J, Li Y, Lei J & Zhou Q (2020b) Structural insight into the substrate recognition and transport mechanism of the human LAT2-4F2hc complex. Cell Discov 6: 82

Yang M, Livnat Levanon N, Acar B, Aykac Fas B, Masrati G, Rose J, Ben-Tal N, Haliloglu T, Zhao Y & Lewinson O (2018) Single-molecule probing of the conformational homogeneity of the ABC transporter BtuCD. Nat Chem Biol 14: 715–722

Zhao Y, Terry D, Shi L, Weinstein H, Blanchard SC & Javitch JA (2010) Single-molecule dynamics of gating in a neurotransmitter transporter homologue. Nature 465: 188–193

Zhao Y, Terry DS, Shi L, Quick M, Weinstein H, Blanchard SC & Javitch JA (2011) Substrate-modulated gating dynamics in a Na+-coupled neurotransmitter transporter homologue. Nature 474: 109–113

Zheng SQ, Palovcak E, Armache J-P, Verba KA, Cheng Y & Agard DA (2017) MotionCor2: anisotropic correction of beam-induced motion for improved cryo-electron microscopy. Nat Methods 14: 331–332

